# TRPV4 is expressed by enteric glia and muscularis macrophages of the colon but does not play a prominent role in colonic motility

**DOI:** 10.1101/2024.01.09.574831

**Authors:** Pradeep Rajasekhar, Simona E Carbone, Stuart T Johnston, Cameron J Nowell, Lukasz Wiklendt, Edmund J Crampin, Yinghan She, Jesse J DiCello, Ayame Saito, Luke Sorensen, Thanh Nguyen, Kevin MC Lee, John A Hamilton, Sebastian K King, Emily M Eriksson, Nick J Spencer, Brian D Gulbransen, Nicholas A Veldhuis, Daniel P Poole

## Abstract

**Background:** Mechanosensation is an important trigger of physiological processes in the gastrointestinal tract. Aberrant responses to mechanical input are associated with digestive disorders, including visceral hypersensitivity. Transient Receptor Potential Vanilloid 4 (TRPV4) is a mechanosensory ion channel with proposed roles in visceral afferent signaling, intestinal inflammation, and gut motility. While TRPV4 is a potential therapeutic target for digestive disease, current mechanistic understanding of how TRPV4 may influence gut function is limited by inconsistent reports of TRPV4 expression and distribution.

**Methods:** In this study we profiled functional expression of TRPV4 using Ca^2+^ imaging of wholemount preparations of the mouse, monkey, and human intestine in combination with immunofluorescent labeling for established cellular markers. The involvement of TRPV4 in colonic motility was assessed *in vitro* using videomapping and contraction assays.

**Results:** The TRPV4 agonist GSK1016790A evoked Ca^2+^ signaling in muscularis macrophages, enteric glia, and endothelial cells. TRPV4 specificity was confirmed using TRPV4 KO mouse tissue or antagonist pre-treatment. Calcium responses were not detected in other cell types required for neuromuscular signaling including enteric neurons, interstitial cells of Cajal, PDGFRα+ cells, and intestinal smooth muscle. TRPV4 activation led to rapid Ca^2+^ responses by a subpopulation of glial cells, followed by sustained Ca^2+^ signaling throughout the enteric glial network. Propagation of these waves was suppressed by inhibition of gap junctions or Ca^2+^ release from intracellular stores. Coordinated glial signaling in response to GSK1016790A was also disrupted in acute TNBS colitis. The involvement of TRPV4 in the initiation and propagation of colonic motility patterns was examined *in vitro*.

**Conclusions:** We reveal a previously unappreciated role for TRPV4 in the initiation of distension-evoked colonic motility. These observations provide new insights into the functional role of TRPV4 activation in the gut, with important implications for how TRPV4 may influence critical processes including inflammatory signaling and motility.

**Summary:**

- TRPV4 is expressed by equivalent cell types in the rodent and primate (monkey and human) colon. This mechanosensitive ion channel has proposed roles in inflammation, visceral afferent signaling, and colonic motility.
- New analysis methods were developed to examine cellular communication in the enteric glial network. This approach revealed new insights into inflammation-associated changes in glial connectivity.
- New roles for TRPV4 in transduction of distension-evoked responses in the colon and colonic motility were identified.

**Key findings:** We have defined the cell types that functionally express TRPV4 in the gut wall. These include enteric glia, endothelia of blood and lymphatic vessels, mMac, and extrinsic afferent nerves. TRPV4- dependent Ca^2+^ signaling was not detected in enteric neurons, PDGFRα cells, interstitial cells of Cajal and smooth muscle cells, which are important drivers of gut motility. These observations align with our experimental evidence for limited involvement of TRPV4 in neuromuscular transmission and propagating colonic motility.

*New and Noteworthy:*

- Novel cellular sites of functional TRPV4 expression in the GI tract were identified and compared across multiple vertebrate species. New analytical approaches to characterize enteric glial communication in a spatiotemporal manner were developed.
- A supporting role for TRPV4 in the initiation of propagating colonic contractions in response to distension was demonstrated. Potential mechanisms that contribute to TRPV4-mediated effects on GI function were identified.
- TRPV4-dependent activity in enteric glia is enhanced in inflammation, consistent with current evidence for inflammation-associated sensitization of TRPV4 on visceral afferents and a major role in mechanically evoked nociceptive signaling.
- Pair correlation analysis was used to examine spatial connectivity of Ca^2+^ signaling, enabling demonstration of dysregulated glial communication in acute inflammation.

## Background

Distension of the gut wall is an important initiator of the coordinated motor patterns that are essential for effective propulsion of content along the gastrointestinal (GI) tract [1, 2]. Recently, there have been significant advances in defining the molecular basis and circuitry of GI mechanosensation and a greater appreciation that a diverse range of cell types are mechanosensitive. Mechanotransducing ion channels including those of the Piezo and Transient Receptor Potential (TRP) families are proposed to be central to this process. For example, roles in intestinal motility have recently been demonstrated for Piezo2 expressed by enterochromaffin cells or extrinsic sensory neurons [2, 3]. The non-selective cation channel Transient Receptor Potential Vanilloid 4 (TRPV4) is another candidate that has been associated with visceral mechanosensation [4, 5]. TRPV4 gating is promoted by diverse modalities including mechanical shear stress, hypo-osmotic changes, lipids and their metabolites, and through coupling to G protein-coupled receptors (GPCRs) and to other ion channels [6, 7, 8, 9]. TRPV4 is therefore able to detect and respond to physiologically important stimuli in the gut as well as to local environmental changes commonly associated with digestive diseases and disorders. Although established and emerging roles for TRPV4 in the GI tract have been identified [10], mechanistic understanding of how these are mediated is limited by current knowledge of where TRPV4 is functionally expressed in the gut wall and how TRPV4 influences cell activity.

TRPV4 is a potential therapeutic target for digestive disease due to its involvement in visceral pain and inflammation. Elevated TRPV4 expression and activity is associated with clinical and experimental inflammatory bowel disease [11, 12]. TRPV4 activation may initiate or promote GI inflammation through immune-derived signaling, neurogenic mechanisms, and disruption of epithelial and endothelial barriers [11, 12, 13]. TRPV4 is also an important driver of mechanically evoked visceral afferent signaling and is associated with post-inflammatory visceral hypersensitivity [4, 5, 14, 15, 16]. TRPV4 expression is upregulated in irritable bowel syndrome (IBS) [11, 12, 17] and levels of endogenous lipid activators of TRPV4 correlate with reported abdominal pain scoring by IBS patients [14]. Furthermore, TRPV4 expressed by sensory neurons is sensitized following exposure to intestinal biopsy supernatants from IBS patients [18] or by prior activation of pronociceptive and proinflammatory GPCRs [15, 16]. Although, these observations support therapeutic targeting of TRPV4 to suppress both intestinal inflammation and visceral hypersensitivity, it remains unclear how TRPV4 inhibition may impact other fundamental gut processes, such as motility. Further insight is therefore required to delineate where and how TRPV4 signaling contributes to GI physiology.

TRPV4 can either inhibit or promote movement of gut contents, mediated through neurogenic and non-neurogenic mechanisms, respectively [17, 19]. Functional expression of TRPV4 by enteric neurons of the mouse, guinea pig, and human intestine has been reported [17, 18, 20], consistent with potential roles in the control of intestinal motility and secretomotor function. In contrast, Luo *et al*. (2018) [19] examined expression using a TRPV4-eGFP reporter mouse and reported that TRPV4 in the external muscle of the intestine was exclusively expressed by resident muscularis macrophages (mMac) and not by other cell types. Selective activation of TRPV4 on mMac had prokinetic effects on GI motility through a PGE2-dependent mechanism. However, it is noted that TRPV4 expression was not detected in cell types where it is abundant and functionally important, such as vascular endothelia [21].

In this study, we aimed to address the key knowledge gaps and conflicting information in the literature using combined Ca^2+^ imaging and immunofluorescent labeling of colonic wholemount preparations. This approach was supported by pharmacological and genetic tools which enabled highly targeted and specific interrogation of the location and identity of TRPV4-positive cells. The spatiotemporal dynamics of TRPV4 signaling in glial cell populations were defined using correlative image analysis and mathematical modelling. The functional role of TRPV4 in motility was determined by measuring neurogenic contractions and complex motor patterns of the isolated mouse colon.

Imaging studies confirmed functional TRPV4 expression by mMac and revealed TRPV4 agonist evoked Ca^2+^ signaling in a subset of enteric glial cells and by lymphatic and vascular endothelial cells. Acute colitis was associated with disruption of coordinated TRPV4 signaling in the enteric glial network and suppressed signaling in lymphatics and mMac. TRPV4 activation or inhibition did not influence neuromuscular transmission or promote colonic contractions and was not required for distension-evoked colonic motility. However, a novel role for TRPV4 in the initiation of distension-evoked propagating colonic motor patterns was identified. These findings challenge current understanding of how TRPV4 contributes to gut motility and are consistent with a role for TRPV4 as a facilitator, rather than a primary mechanosensor or driver of colonic motility.

## Materials and Methods

### Mice

The following mouse strains were used: C57Bl/6J, TRPV4^-/-^ [22], Wnt1-GCaMP3 [23], *Csf1r*-eGFP (MacGreen, [24]; RRID:IMSR_JAX:018549), and CGRPα-mCherry [25] (6-14 weeks, male and female). All procedures involving mice were approved by animal ethics committees of the Monash Institute of Pharmaceutical Sciences (12199, 14816) and Flinders University (933-16). All mice were housed under a 12 h light/dark cycle, with controlled temperature (24°C) and humidity, and free access to food and water.

### Human tissues

Healthy tissue from corrective surgery for anorectal malformation was obtained from the Murdoch Children’s Research Institute at The Royal Children’s Hospital (RCH, Melbourne). Consent was obtained from guardians and collection was approved by the Human Research Ethics Committee at RCH (38262). Patient data are provided in **Table S1**.

### Monkey tissues

Rectal tissues from 6 healthy pigtail macaques (*Macaca nemestrina*) were obtained from an unrelated HIV vaccine study (specific details related to this study and animal welfare are provided in [26]; Australian Animal Health Laboratory protocol approval number 1887). Animals received vaccines but were not infected. Tissues were transported in complete RPMI-1640 (Sigma-Aldrich) containing 25% fetal calf serum (Gibco) before transfer to Krebs buffer and preparation of wholemounts for Ca^2+^ imaging studies.

### Ca^2+^ imaging of wholemount preparations of the colon

Myenteric and submucosal wholemount preparations were incubated with Fluo8-AM Ca^2+^ indicator dye (6 μM in loading solution: 4 mM probenecid, 0.02% pluronic F-127, 0.1% cremophor-EL and 0.5% BSA in L-15 medium, pH 7.4) in a humidified incubator (40 min, 37°C), then recovered in Krebs buffer for 10 min. Tissue imaging was performed using a Leica TCS SP8 multiphoton system with a HC PLAN APO 0.95 NA 25x water immersion objective. The microscope stage and buffer were maintained at 37°C. The intensity of Fluo8 fluorescence, which is indicative of changes in intracellular Ca^2+^ concentration ([Ca^2+^]_i_), was detected at 488 nm excitation and 530±20 nm emission. Images were recorded at 1 frame per 0.86 s, 1024 x 1024-pixel resolution and scan frequency of 600 Hz. Equivalent settings were used to detect changes to the fluorescence intensity of the Ca^2+^ biosensor GCaMP3. Drugs were applied directly to the bath, close to the field of view. Effective mixing was confirmed by rapid Ca^2+^ responses by cells to ATP.

A series of experiments used higher speed imaging of myenteric neuron activity or tissue from CGRPα-mCherry mice to selectively measure visceral afferent activity. In these cases, wholemount preparations were loaded with Fluo-4 AM Ca^2+^ indicator dye (4 μM) in Krebs buffer. Images were acquired using a Nikon Eclipse Ci-S widefield upright microscope with 20x water immersion objective at a resolution of 512 x 512 pixels and a speed of 70 Hz [25].

### Drugs

Pharmacological tool compounds and other reagents used in this study are summarized in **Table S2**.

### Immunofluorescence

Following Ca^2+^ imaging, tissues and cells were fixed and labeled using indirect immunofluorescence as described [27]. Details of the selective cellular markers and the antibodies used to detect these are provided in **Table S3**. Preparations were imaged using a TCS-SP8 confocal system (20x HC PL APO NA 1.33) or TCS-SP8 Lightning confocal system (20x HC PL APO NA 0.88 or 40x HC PL APO NA 1.1 water immersion).

### Image analysis

The timeseries images were registered in FIJI (ImageJ v1.52c) and aligned with the corresponding immunolabeled image to identify specific cell types. Regions of interest (ROIs) corresponding to individual cells were then identified, enabling extraction of mean intensity values for Ca^2+^ responses and the cell coordinates for further analysis. A detailed description of the methods used is provided in the ***supplementary methods***.

### Analysis of spatial connectivity of enteric glia

Pair correlation function (PCF) analysis was used to determine coordination of glial responses. Here PCF is a measure of how closely the response of glia at different locations is correlated, where distance between locations is measured as a path length within myenteric ganglia. PCFs were calculated from glial Ca^2+^ responses, normalized using the ganglionic area within the myenteric plexus and their locations within the ganglia. The distances between pairs of glial cells were calculated via a computational meshing procedure and PCF calculated from the corresponding adjacency matrix [28, 29]. PCF values were determined for glia that responded within 30s and 60s after drug or vehicle addition. PCF was normalized such that values >1 identified coordinated glial activity. In contrast, uncoordinated or independently active glia generated PCF values ∼1 for all pair distances and times (**Fig. S1**). The following PCF values were used to indicate correlations: strong correlation >1.4; moderate 1.2-1.4; weak 1-1.2, and negative <1.

### Macrophage depletion

Macrophages were depleted using an anti-CSFR1 mAb as described ([30], ***supplementary methods***).

### TNBS colitis

Acute TNBS colitis was established according to the presensitization model outlined by Wirtz et al. 2017 [31] (***supplementary methods***).

### Colonic contraction and motility assays

Changes in the tension of circular muscle strips of the mouse proximal colon were measured as described [32]. TRPV4-mediated effects on contractile activity and electrically evoked contractions were quantified. Propagating motility was examined in the isolated, intact mouse colon by video imaging and spatiotemporal mapping [33]. The time interval between colonic motor contractions (CMCs), CMC velocity and resting gut diameter between contractions were determined from diameter maps (DMaps). The involvement of TRPV4 in resting colonic diameter prior to contraction onset and the relative rate of tissue contraction was assessed using custom analysis of DMaps. Mean rate of contractions (MRC) was defined as follows: *change in gut diameter at the onset to the peak of the contraction in mm/ time taken to reach the peak in seconds*. A detailed overview of these analyses is provided in the ***supplementary methods*** and **Fig. S2**.

**Figure 1.**
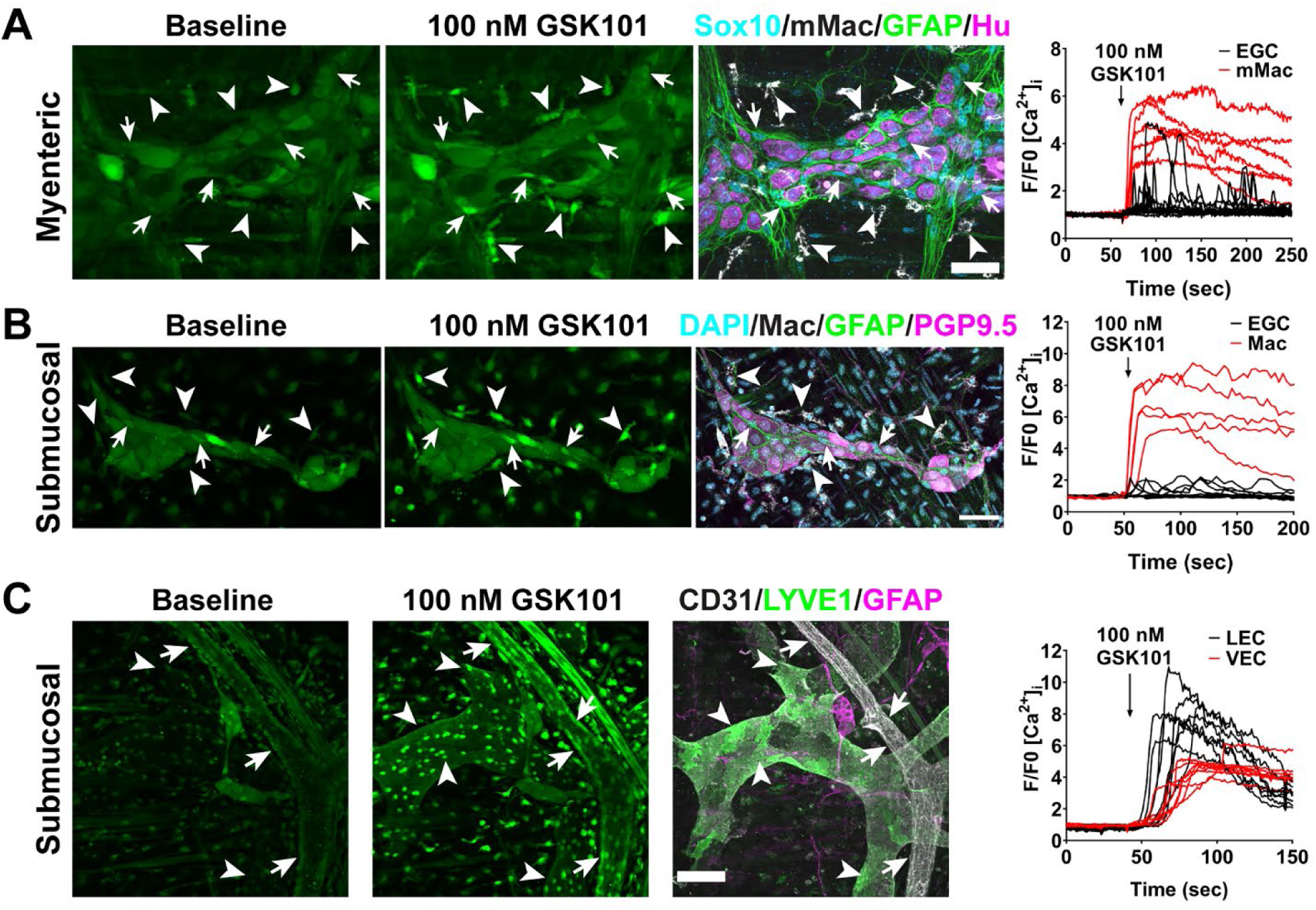
TRPV4 is functionally expressed by enteric glia, macrophages, and endothelial cells of the mouse colon. **A.** Enteric glia (arrows) and mMac (arrowheads) in the myenteric region exhibited increased [Ca^2+^]i in response to the TRPV4 agonist GSK101 (100 nM). Immunostaining confirmed the identity of glial cells (Sox10+: cyan, GFAP: green), mMac (white), and neurons (Hu+: magenta). Scale = 50 µm. **B.** Responses to GSK101 were also detected in glial cells (arrows) and macrophages (arrowheads) associated with submucosal ganglia. Scale = 50 µm. **C.** GSK101 evoked sustained Ca^2+^ signals in vascular (arrows, CD31+: white) and lymphatic (arrowheads, LYVE1+: green) endothelial cells. Each trace represents the response by an individual cell. Scale = 100 µm.

**Figure 2.**
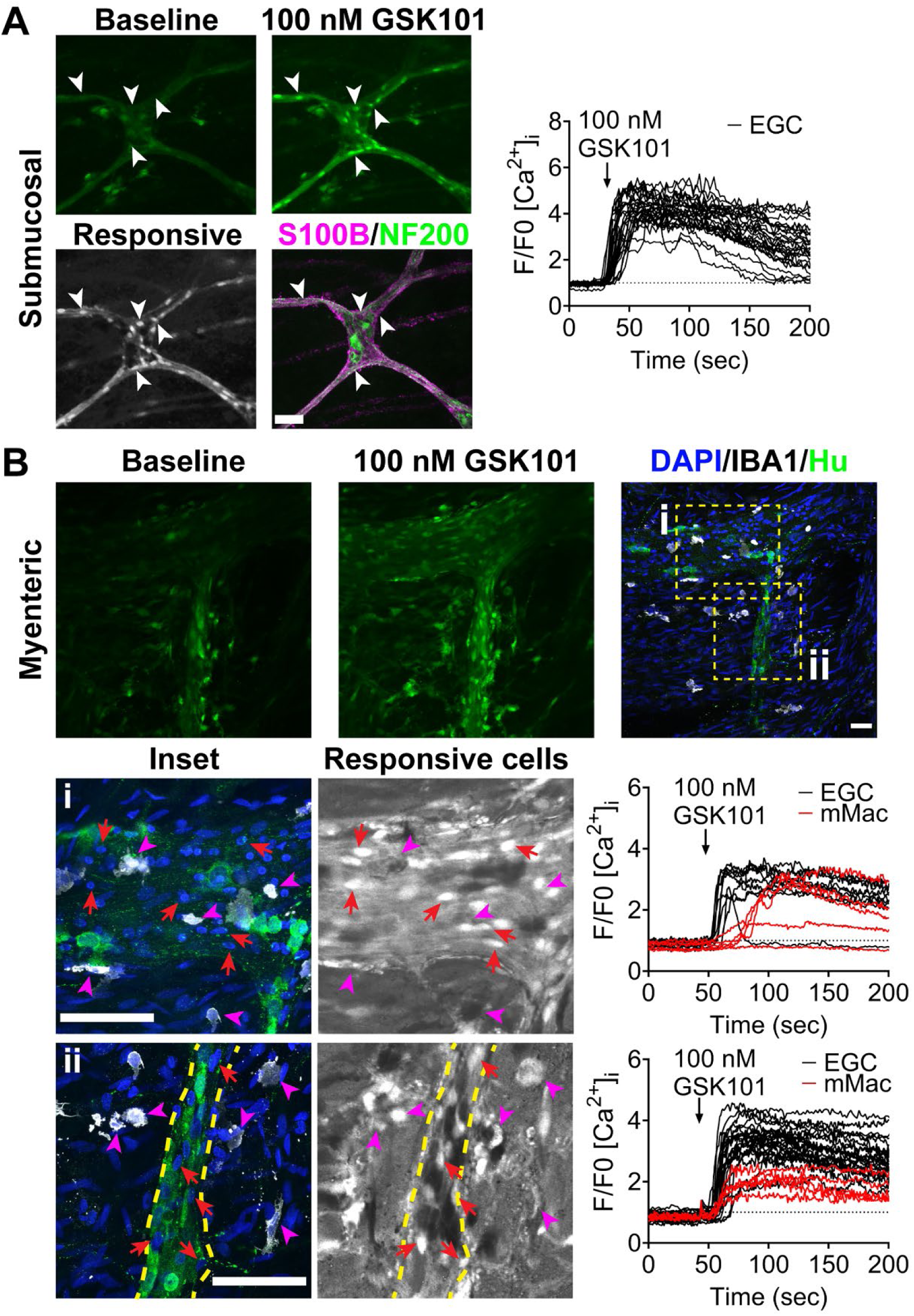
TRPV4 is functionally expressed by enteric glia and macrophages in the human colon. **A.** GSK101 (100 nM) induced sustained Ca^2+^ responses in submucosal glia (arrowheads; S100B+: magenta) of the human colon. **B.** Equivalent responses were detected in the myenteric region. Insets **i** & **ii** show responses by myenteric glia (red arrows) and macrophages (magenta arrowheads; scale: 20 µm). Enteric glia were identified as non-neuronal cells (DAPI: blue) that were closely associated with neurons (Hu: green) within the ganglia. Macrophages were defined as IBA1+ cells (white). The grayscale images highlight responsive cells in white against a dark background (image normalized to baseline, F/F0). Scale = 50 µm.

### Statistical Analysis

Data are expressed as mean ± 95% CI or SD. Statistical tests, replicates, and post-hoc analyses for each experiment are as indicated in respective figure legends.

## Results

### TRPV4 is expressed by macrophages and glia in the mouse colon

Calcium imaging of myenteric wholemount preparations was used to establish the precise distribution of cells responsive to TRPV4 agonists in the mouse colon. The highly selective TRPV4 agonist GSK101 (100 nM) evoked robust Ca^2+^ responses by distinct cell types, which were identified by immunofluorescent labeling for established markers (**Table S3**) and through use of macrophage reporter mice. GSK101 stimulated an immediate and sustained elevation in intracellular Ca^2+^ in all mMac examined (96.6 ± 2.4% of total mMac population, mean ± 95% C.I.; N= 222 cells, n=9 mice) (**Fig. 1A, Video S1**). Responses were also observed in blind-ended initial lymphatic vessels, and blood vessels (*not shown*). Ca^2+^ responses by glia located in myenteric ganglia were transient and oscillatory, with a delayed onset relative to other cell types (61.5±10.4% of total glial population, N=1804 cells, n=13 mice: **Fig. 1A**). In contrast, no direct responses by myenteric neurons, interstitial cells of Cajal (ICC), PDGFRα cells (Fig. S3), and intestinal smooth muscle cells were detected (*not shown*). Responses to GSK101 were retained in the presence of tetrodotoxin (TTX, 1 µM).

**Figure 3.**
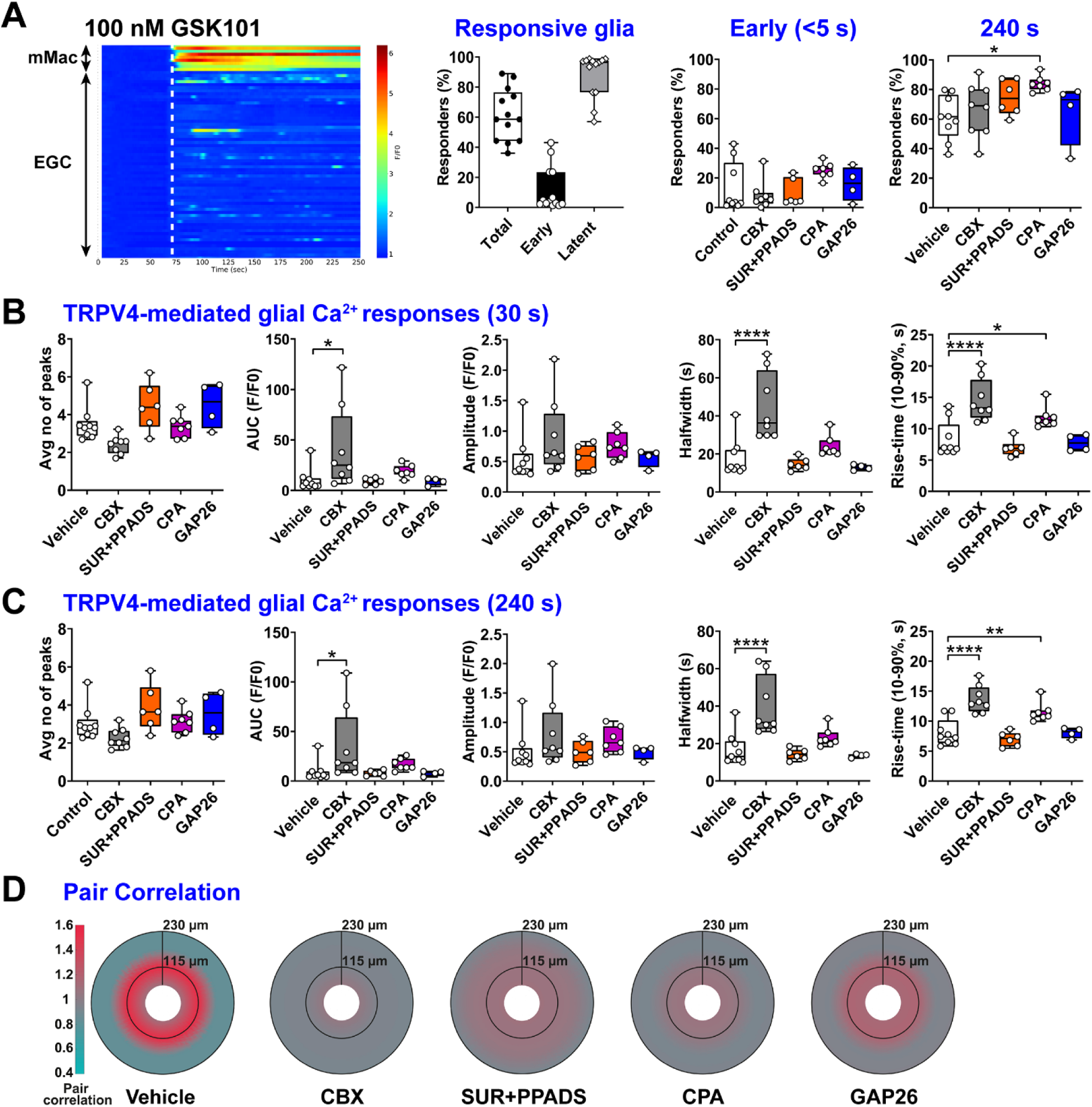
TRPV4-mediated activation of the myenteric glial network of the mouse colon is mediated by gap junctions, purinergic activity, and release of Ca^2+^ from intracellular stores. **A.** Heatmap demonstrating the oscillatory nature and varying kinetics of Ca^2+^ responses by glia to TRPV4 activation (100 nM GSK101, white dotted line). Responses by mMac to the same stimulus were sustained. A distinct subset of the GSK101-responsive glial population was identified that exhibited rapid (‘early’) elevations in Ca^2+^ within 5s of GSK101 application. The mechanism of GSK101-induced Ca^2+^ activity within the glial network was investigated by pharmacological inhibition of gap junctions (50 µM carbenoxolone: CBX), purinoceptors (50 µM suramin (SUR) + 50 µM PPADS), connexin43 hemichannels (50 µM GAP26), and depletion of intracellular Ca^2+^ from ER stores (30 µM cyclopiazonic acid: CPA). None of the pharmacological treatments significantly affected early responding glial populations (<5 s). However, there was a significant increase in the percentage of responding glia following CPA pre-treatment when assessed at 240 s. The kinetics of the Ca^2+^ responses were compared at t= 30 s **(B)** and 240 s **(C)**. CBX pre-treatment resulted in a significant increase in AUC, halfwidth and rise-time (10-90%) for cells responding both within 30 s and 240 s post GSK101 addition, compared to vehicle. In the presence of CPA, there was an increase in rise-time for cells responding at both time points. (Vehicle: 1045 cells, N=9; CBX: 936 cells, N=8; CPA: 917 cells, N=7; Gap26: 503 cells, N=4; SUR+PPADS: 704 cells, N=6). **(D)** Pair correlation function (PCF) analysis of responses at t= 30 s for cells separated by 115 µm revealed smaller PCF values across all treatments compared to vehicle (PCF: Veh= 1.54, CBX= 0.93, SUR+PPADS= 1.02, CPA= 1.03, Gap26= 1.17). This indicates that the correlated glial activity was dependent on gap junctions, purinergic activity and intracellular Ca^2+^ stores. Data were analyzed by one-way ANOVA with Dunnett’s post hoc test, * P<0.05, ** P<0.01, **** P<0.0001. Data are expressed as mean ± 95% C.I. The horizontal line in each graph represents the mean, and whiskers denote the minimum and maximum values.

TRPV4 function was also examined in submucosal wholemounts. Addition of GSK101 resulted in elevated intracellular Ca^2+^ in macrophages, a subset of glia (18/52 cells, n=4 mice) (**Fig. 1B**), blood and lymphatic vessels (**Fig. 1C**), and extrinsic afferent nerve fibers (**Fig. S4**, **supplemental data**). Responses by lymphatic endothelia and macrophages were immediate and sustained (traces in **Fig. 1B, C**), whereas blood vessels exhibited rapid oscillatory fluctuations in Ca^2+^ (*not shown*).

**Figure 4.**
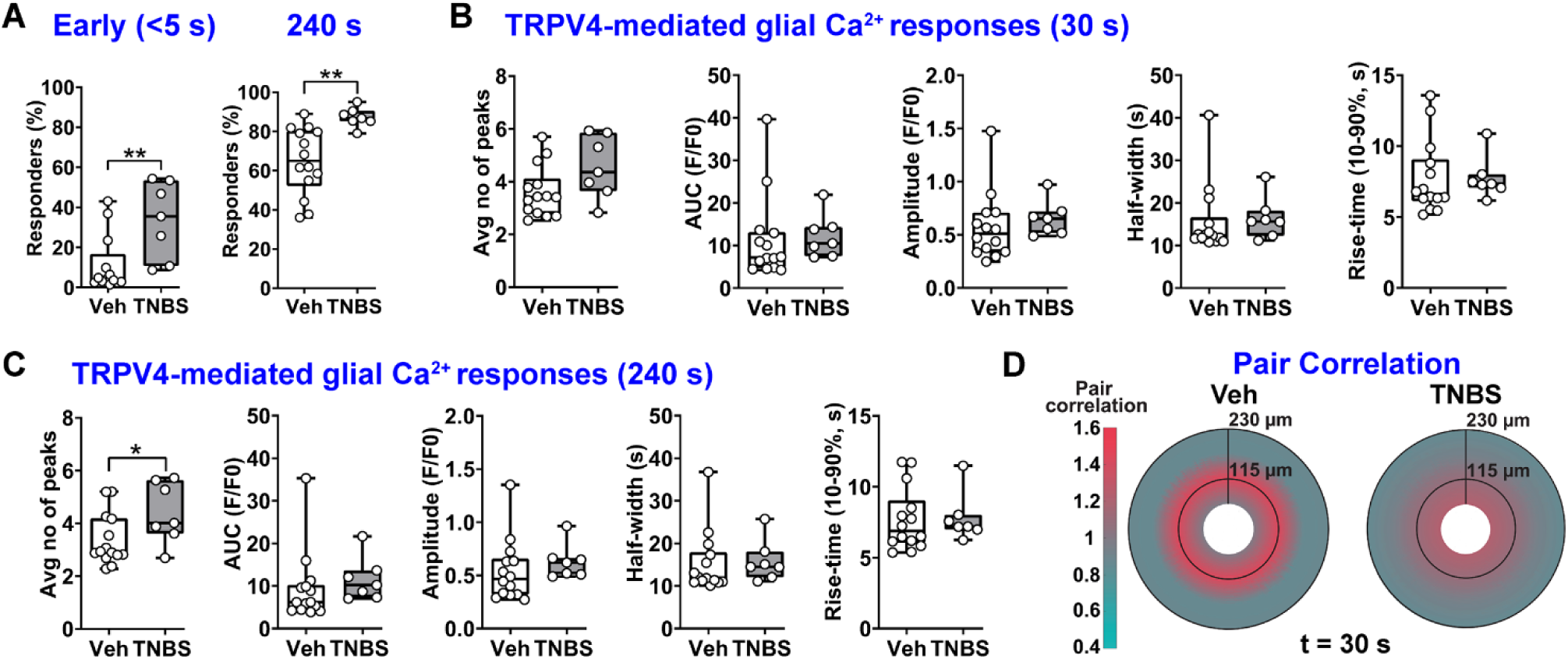
TRPV4-mediated glial activity is sensitized in acute TNBS-induced colitis and is associated with a reduction in correlated Ca^2+^ activity. **A.** The percentage of GSK101 (100 nM)-responsive glia was increased in acutely inflamed tissues at both the early and late time points. **B.** GSK101-induced Ca^2+^ kinetics were not affected when assessed at 30 s post drug addition. However, there was an increase in the average number of peaks at 240 s **(C)**. **D.** TRPV4-mediated Ca^2+^ responses in the glial network were less coordinated for cells within a 115 µm distance in inflamed tissues (PCF= 1.05) compared to the healthy control (PCF= 1.44). (Vehicle: 1260 cells, N=14; TNBS: 598 cells, N=7; one-way ANOVA with Dunnett’s post hoc test, * P<0.05, ** P<0.01. Data are expressed as mean ± 95% C.I. The horizontal line in the graph represents the mean, and whiskers denote the minimum and maximum values.

The selectivity of GSK101 for TRPV4 was confirmed by pharmacological and genetic approaches. All Ca^2+^ responses to GSK101 were abolished by the TRPV4 blockers HC067047 (HC, 10 µM, n=7; **Fig. S5A**) and GSK2193874 (GSK219, 1 µM, n=3) (**Fig. S5B**), and absent in TRPV4^-/-^ tissues (n=9, **Fig. S5C**). Effective dye loading and cell viability were verified based on Ca^2+^ responses to ATP (100 µM). These observations demonstrate the selectivity of GSK101 for TRPV4 and support its use as a tool compound to probe TRPV4 function by Ca^2+^ imaging.

Identification of where TRPV4 is functionally expressed in tissues may be confounded by indirect activation of cells following release of signaling mediators and by the high degree of intercellular connectivity. To address this issue, functional expression of TRPV4 was confirmed in relevant primary cells and commonly used cell lines. TRPV4 signaling was directly assessed in cultured primary human and rodent macrophages, mouse enteric glia, and human dermal lymphatic endothelial cells, and related cell lines (**Figs S6 & S7**). Data from these studies confirmed the observations made in tissue preparations.

These findings demonstrate that TRPV4 is functionally expressed by mMac and glia, two cell types with emerging roles in gut motility [34, 35]. Notably, TRPV4 was not expressed by myenteric neurons, ICC, and PDGFRα^+^ cells, which are important cell types directly involved in neuromuscular transmission and the control of motility [35]. TRPV4 was also prominently expressed by vascular and lymphatic endothelia, consistent with involvement in the control of fluid balance and proposed roles in edema and inflammation [8, 36, 37].

### TRPV4 is expressed by macrophages and glia in the submucosal and myenteric regions of the human and monkey intestine

Differences in the expression, distribution, and function of cell surface receptors between species can have significant implications for interpretation of their function and for future therapeutic targeting. To address this potential limitation, we examined the functional distribution of TRPV4 in wholemount preparations of the human colon (**Fig. 2A, B, Video S2**) and monkey rectum (**Fig. S8**) using Ca^2+^ imaging. GSK101 (100 nM) evoked rapid and robust responses by myenteric glia and mMac in both human and monkey tissue (<u>human</u>-glia: 92.6 ± 19.7%, 768 cells; mMac: 94.8 ± 6.6%, 55 cells, n=5; <u>monkey</u>-glia: 83.3 ± 15.1%, 242 cells, n=5; mMac: 100%, 58 cells, n=6). There was no evidence for neuronal activation in these preparations, consistent with observations from the mouse colon.

GSK101 also evoked rapid and sustained increases in Ca^2+^ in glial cells in submucosal wholemounts of the human colon (**Fig. 2A**). All responses were abolished by HC067 (n=4) (**Fig. S5D**).

These data demonstrate that the functional distribution of TRPV4 is consistent across species. Further, these observations support the use of the mouse colon for assessment of the role of TRPV4 in physiological processes, such as gut motility.

### TRPV4 promotes activation of the glial network through gap junction coupling

TRPV4 activation of astrocytes induces intercellular Ca^2+^ waves which are mediated via gap junctions, ATP release [38], and IP_3_R-dependent Ca^2+^ induced Ca^2+^ release [39]. Similar mechanisms promote activation and propagation of Ca^2+^ waves in glia [40, 41, 42]. We used pharmacological blockers of these pathways to examine the underlying mechanisms that drive TRPV4-evoked Ca^2+^ oscillations within the glial network.

Glial Ca^2+^ responses were categorized as early and late responders (<u>early</u>: initial response within 5s, 11.9% ± 8.8, n=13; <u>late</u>: >5s, 88% ±14.4, n=13) (**Fig. 3A**). Equivalent data were obtained using tissues from Wnt1-GCaMP3 mice, which enabled examination of Ca^2+^ signaling in the absence of probenecid, which can inhibit neuronal pannexin channels [43] (**early:** 13.5% +/- 7.2, **total responders:** 76.6% +/- 5.4, n=3).

Neuron-glia interactions are involved in the generation and maintenance of intercellular Ca^2+^ waves in glia [40, 43]. However, TRPV4-evoked Ca^2+^ oscillations persisted when neurotransmission was blocked by TTX (1 µM) (**Fig. S9**, **Table 1**) (n=591 cells, N=6). Activity was also retained in the presence of the purinoceptor antagonists suramin (50 µM) and PPADS (50 µM) (n=704 cells, N=6) (**Fig. 3A-C, Video S3**), consistent with an ATP-independent mechanism of action. Effective inhibition was confirmed by block of Ca^2+^ responses to exogenous ATP (100 µM). Release of Ca^2+^ from intracellular stores can promote release of mediators which drive cellular communication. The SERCA inhibitor cyclopiazonic acid (CPA; 30 µM) significantly increased the percentage of GSK101-responsive glia and the rise time of GSK101-evoked responses (n=917 cells, N=7) (**Fig. 3A-C**). Ca^2+^ responses by glia to GR64349, an agonist of the G_q_-coupled neurokinin 2 receptor, were inhibited by CPA, confirming effective block of SERCA ([40]; **Video S4**). To assess the role of junctional communication we used the gap junction inhibitor carbenoxolone (CBX) and Gap26, a peptide inhibitor of connexin43 hemichannels [44]. CBX (50 µM) suppressed Ca^2+^ oscillations and revealed a glial subset that exhibited rapid and sustained responses to GSK101 (n= 936 cells, N= 8) (**Fig. 3A-C**, **Fig. S10B**, and **Video S5**). This was reflected by an increased rise-time, AUC, and half-width of the Ca^2+^ response. Gap26 (50 µM) did not affect GSK101-evoked Ca^2+^ signaling under the experimental conditions used (n=503 cells, N=4) (**Fig. 3A-C**). The effects of the different inhibitors on key Ca^2+^ signaling parameters are summarized in **Table 2 and Fig. S10A**.

**Table 1.**
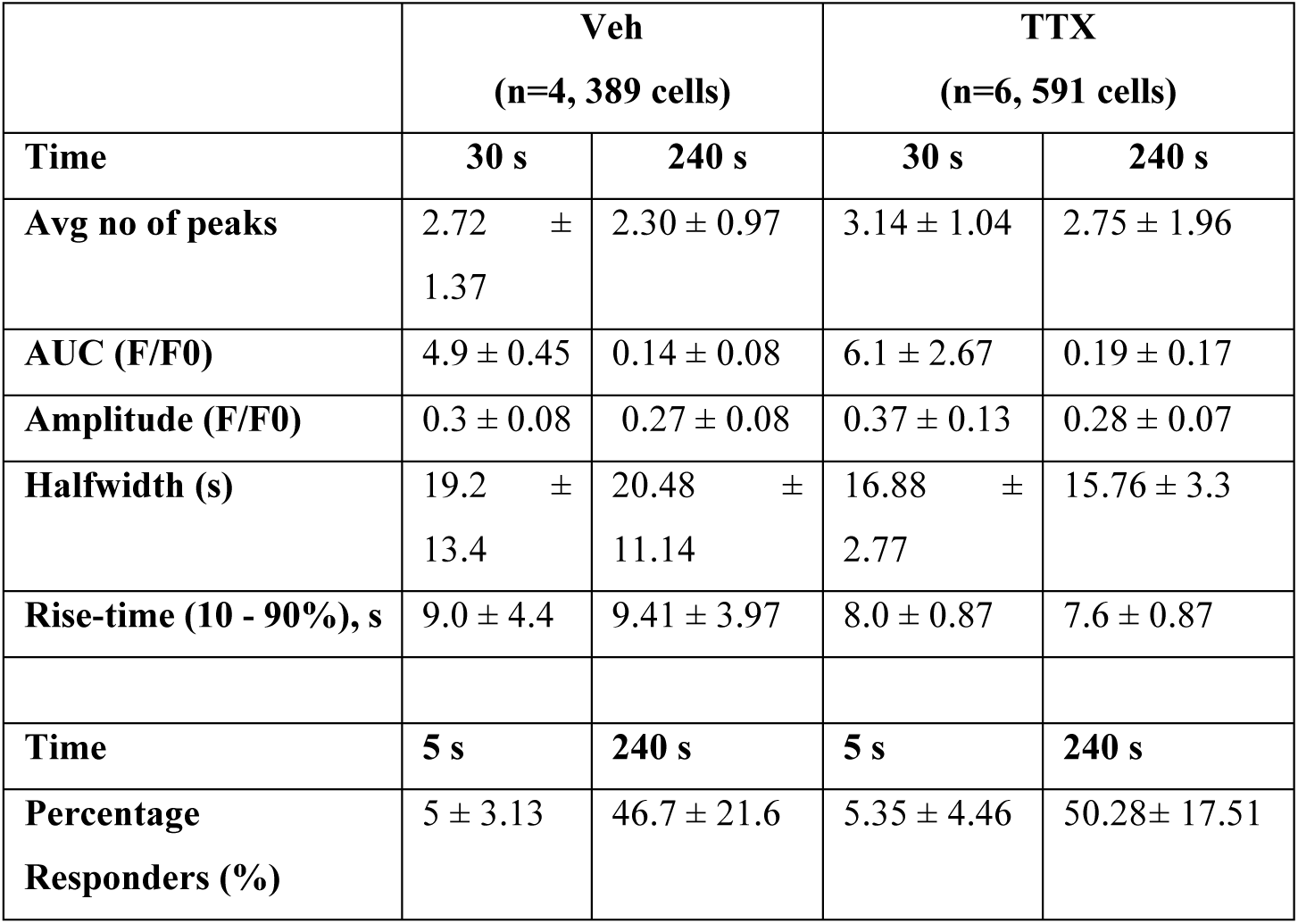
Temporal parameters for TRPV4-mediated glial Ca^2+^ responses in the presence of TTX.

**Table 2.**
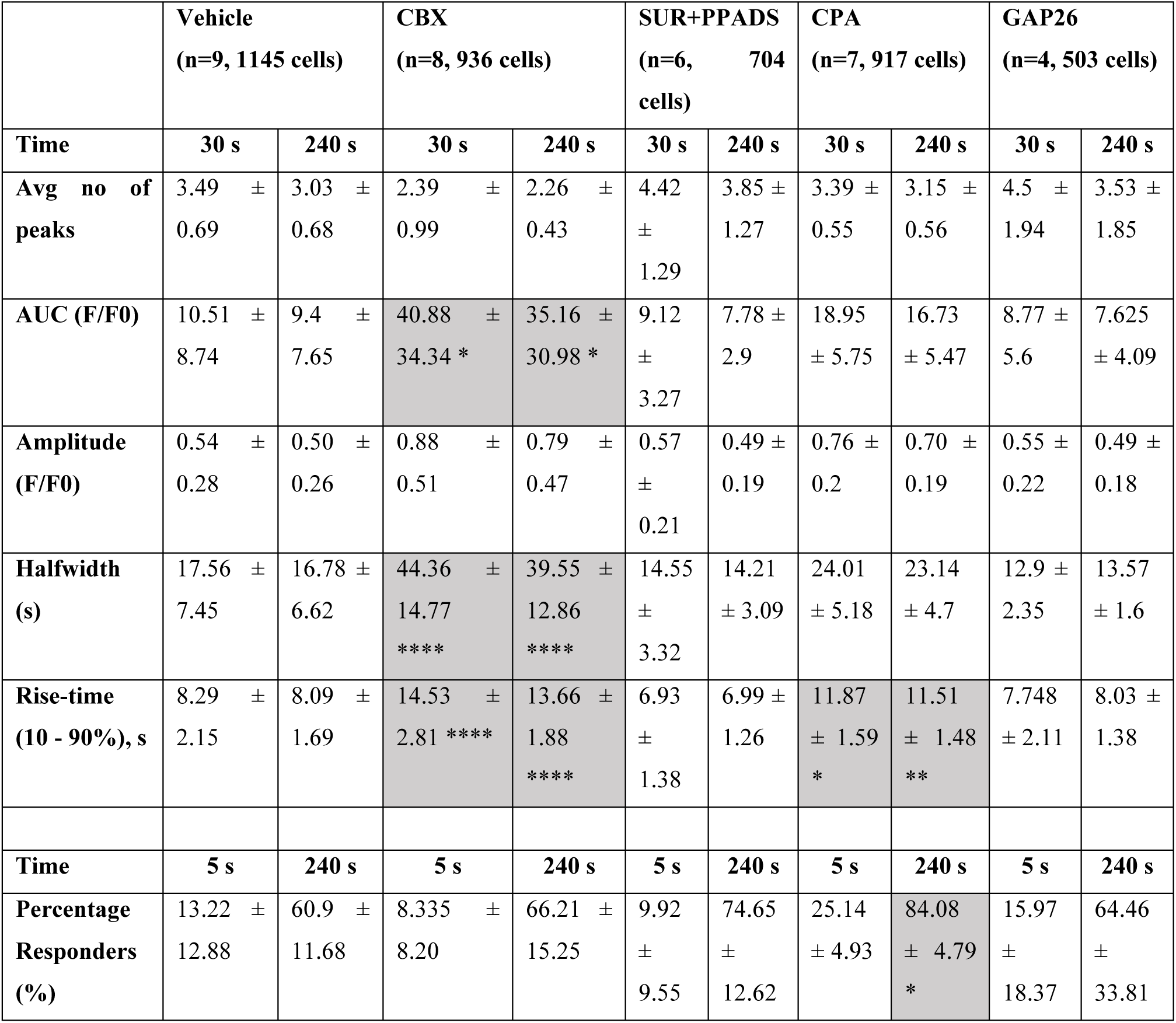
Temporal parameters for TRPV4-mediated glial Ca^2+^ responses in the presence of inhibitors.

**Table 3.** Temporal parameters for TRPV4-mediated glial Ca^2+^ responses in a TNBS-model of colitis.

|  | **Veh**<br>**(n=14, 1924 cells)** |  | **TNBS**<br>**(n=7, 685 cells)** |  |
| --- | --- | --- | --- | --- |
| **Time** | **30 s** | **240 s** | **30 s** | **240 s** |
| **Avg no of peaks** | $3.6 \pm 0.55$ | $3.3 \pm 0.55$ | $4.55 \pm 1.09$ | $4.4 \pm 1.06^*$ |
| **AUC (F/F0)** | $11.28 \pm 5.7$ | $9.35 \pm 4.7$ | $12.14 \pm 4.7$ | $11.52 \pm 4.7$ |
| **Amplitude (F/F0)** | $0.57 \pm 0.18$ | $0.54 \pm 0.16$ | $0.65 \pm 0.15$ | $0.63 \pm 0.15$ |
| **Halfwidth (s)** | $15.6 \pm 4.6$ | $15.3 \pm 4.18$ | $16.4 \pm 4.5$ | $15.8 \pm 4.53$ |
| **Rise-time (10 - 90%), s** | $7.72 \pm 1.5$ | $7.67 \pm 1.27$ | $7.74 \pm 1.3$ | $7.81 \pm 1.58$ |
| **Time** | **5 s** | **240 s** | **5 s** | **240 s** |
| **Percentage Responders (%)** | $11.06 \pm 8.5$ | $64.9 \pm 9.9^{**}$ | $33.58 \pm 17.73$ | $87.2 \pm 4.6^{**}$ |

Exclusive expression of TRPV4 by mMac [19] and functional interaction between mMac and enteric neurons [45] and glia [46] have been reported. It is therefore possible that the sustained glial activity observed results from continued input from closely associated mMac. This potential mechanism was tested by examining glial Ca^2+^ signaling in tissues from animals treated with anti-CSF1R antibodies to deplete macrophages. Responses by glia and by lymphatic and blood vessels were retained in the absence of mMac and in tissues from mice treated with the isotype control (**Fig. S11**). The percentage of GSK101-responsive glia was not significantly different between preparations from mAb (44.7± 19.6%, 1029 cells total) and IgG (62.0±19.2%, 961 cells total) treated mice (n=4 mice per group).

Standard analysis of Ca^2+^ signaling only accounts for temporal features of responses. Furthermore, temporal analysis does not consider the spatial structure of glia within ganglia and how they communicate as part of a network. To address this limitation, pair correlation analysis was used to assess the spatiotemporal dynamics of Ca^2+^ signaling in the glial network (***supplementary methods***, **Fig. S1**). PCFs [29] were used to determine whether TRPV4-induced glial activity was correlated across spatial regions, and if this was affected by the inhibitors examined. PCF values >1 at short pair distances and at earlier time points are indicative of coordinated glial activity (i.e., activation of one glial cell leads to activation of surrounding glia). In contrast, random glial activity is defined by a PCF value =1 at all pair distances and times. Analysis of PCFs at different time points and distances, and relevant controls are provided in ***supplemental data***.

The spatiotemporal dynamics of glial responses as they propagated throughout the network were assessed at 30s post-GSK101 treatment. The PCF values at 115 µm demonstrated a reduction in correlated glial activity with CBX (0.94± 0.17, mean±95% CI), CPA (1.04± 0.05), GAP26 (1.17± 0.17) and suramin + PPADS (1.07± 0.04) compared to vehicle control (1.54±0.91) (**Fig. 3D**). Equivalent analysis of TRPV4-dependent glial signaling in the presence of TTX (**Fig. S9**, **Table 1**) revealed that although activity was retained, the responses at a distance of 115µm (1.03± 0.12) were reduced relative to vehicle control (1.39± 0.72). A summary of PCF values at different distances is provided in **Table S4**. This novel analysis approach reveals that TRPV4-dependent Ca^2+^ responses by glia are modulated via gap junctions, ATP, intracellular Ca^2+^ stores and are reinforced by neuronal activity. We demonstrate that PCF analysis is a valuable tool to reveal changes in glial network activity that are not detectable using standard temporal analyses.

### TRPV4 signaling in the glial network is disrupted in acute inflammation

Intestinal inflammation is associated with increased TRPV4 expression and sensitization of TRPV4 channel activity [10, 12]. Tissues from mice with acute TNBS colitis (3d) were assessed to determine the effects of inflammation on the spatiotemporal dynamics of TRPV4 signaling in individual glial cells and within the glial network. Acute inflammation was associated with a significant increase in the relative percentage of glia that responded to GSK101 within 5s (64.9 ± 9.9% vs. 11.06 ± 8.5%) or at 240s post-addition (87.2 ± 4.6% vs. 33.5 ± 17.7%) (**Fig. 4A, Video S6**). The average number of Ca^2+^ peaks per glial cell was equivalent to control for early responders (<5s) (**Fig. 4B**) but was significantly increased (4.4 ± 1.06) for all responsive glia (240s) (**Fig. 4C**), consistent with increased glial activity. The AUC, amplitude, half-width, and rise-time of responses to GSK101 were not significantly affected by acute inflammation at 30s or 240s (**Table 3**). However, PCF analysis (30s, 115 µm) revealed a disruption of the coordinated activity within the glial network in acutely inflamed tissue (PCF= 1.05± 0.06) relative to controls (PCF= 1.44± 0.77) (**Fig. 4D, Table S4**). These data demonstrate that although inflammation is associated with an increase in the number of responsive glia and an upregulation of glial activity in the inflamed colon, glia within the network respond in an uncoordinated manner. These data are consistent with reports of altered TRPV4 expression or function in acute inflammation.

The effects of acute inflammation on TRPV4-dependent signaling in mMac and LEC were also examined in the same preparations. Acute colitis was associated with a generalized suppression of signaling in both cell types, reflected by significant reductions in the amplitude and area under the curve of responses to GSK101 (**Fig. S12**. These observations contrast with increased responses by glia and demonstrate that disease-associated changes in TRPV4 activity are cell type dependent.

### TRPV4 is not functionally expressed by enteric neurons

TRPV4 is a candidate modulator of intestinal motility based on the mechanosensory properties of this channel. This role is supported by experimental evidence for both inhibitory [17] and prokinetic [19] effects of channel activation, which are mediated through distinct mechanisms. Our initial examination using Ca^2+^ imaging did not detect GSK101-evoked Ca^2+^ transients in enteric neurons. These neurons responded to exogenous ATP, confirming effective dye loading and viability (**Fig. 5A**). TRPV4 function was also measured in neurons from Wnt1-GCaMP3 transgenic mice where expression of the Ca^2+^ biosensor GCaMP3 is restricted to neural crest-derived enteric neurons and glia [23]. GSK101-evoked Ca^2+^ responses were not detected in myenteric neuronal cell bodies in these tissues (923/925 neurons, N=7 mice), although more subtle responses in neurites or varicosities cannot be excluded. Closely associated glia exhibited robust elevations in Ca^2+^ **(Fig. 5B**) including in glial processes. Responses in submucosal ganglia were similarly restricted to glia (**Fig. 5C**). Higher speed imaging was used to account for rapid, transient neuronal responses. No direct activation of myenteric neurons or increase in spontaneous neuronal activity was detected following treatment with GSK101 (**Fig. S13)**. TRPV4 function was examined in cultured myenteric neurons from the mouse colon to determine if phenotypic changes that occur *in vitro* may account for previous observations [17]. Using correlative imaging and more advanced methodology to those in our previous study, we detected no responses to GSK101 by Hu-immunoreactive neurons (69/70 neurons, N=3 independent cultures). In contrast, robust responses by GFAP-and Sox10-positive glia were detected (35/35 glia), including by those that were closely associated with neurons (**Fig. 5D**).

**Figure 5.**
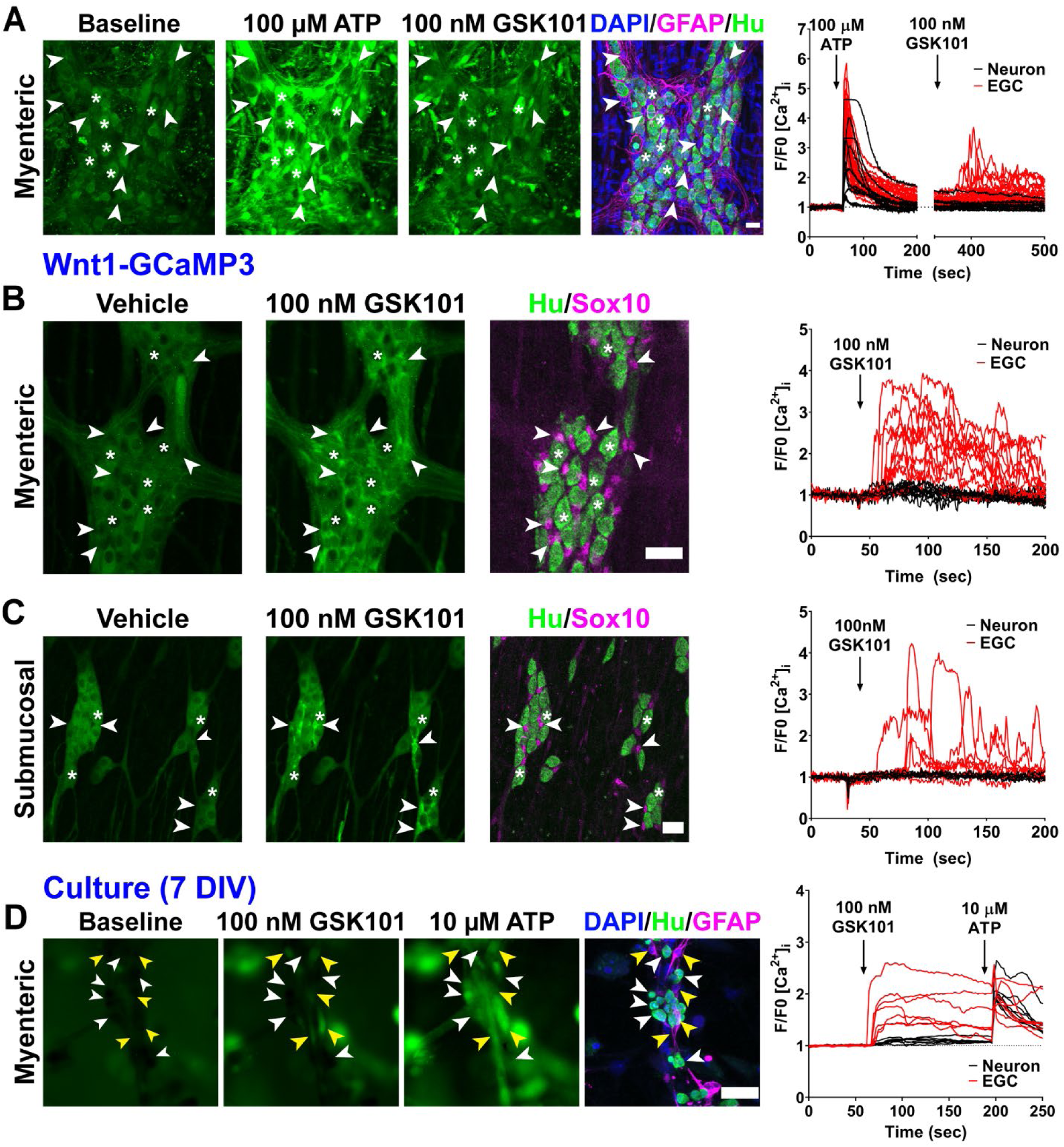
Enteric neurons of the mouse colon do not express functional TRPV4. **A.** Wholemount preparation loaded with Fluo8-AM dye. Myenteric neurons (asterisks; Hu+: green) responded to ATP (100 µM) but were unresponsive to GSK101 (100 nM). In contrast, enteric glia (arrowheads; GFAP+: magenta) responded to both agonists. **B-C.** Wholemount preparation of the Wnt1-GCaMP3 mouse colon. GSK101 evoked Ca^2+^ responses only in enteric glia (arrowheads; Sox10+: magenta) and not enteric neurons (asterisks; Hu+: green) (representative examples from n=5). **D.** Cultured enteric glia (yellow arrowheads; GFAP+: magenta), but not myenteric neurons (white arrowheads; Hu+: green), exhibited increased [Ca^2+^]i in response to GSK101. Both neurons and glia responded to subsequent exposure to ATP (10 µM). In contrast to observations from tissue-based imaging, GSK101-mediated [Ca^2+^]i responses by glia were sustained (n=3 independent cultures). Scale = 40 µm.

**Figure 6.**
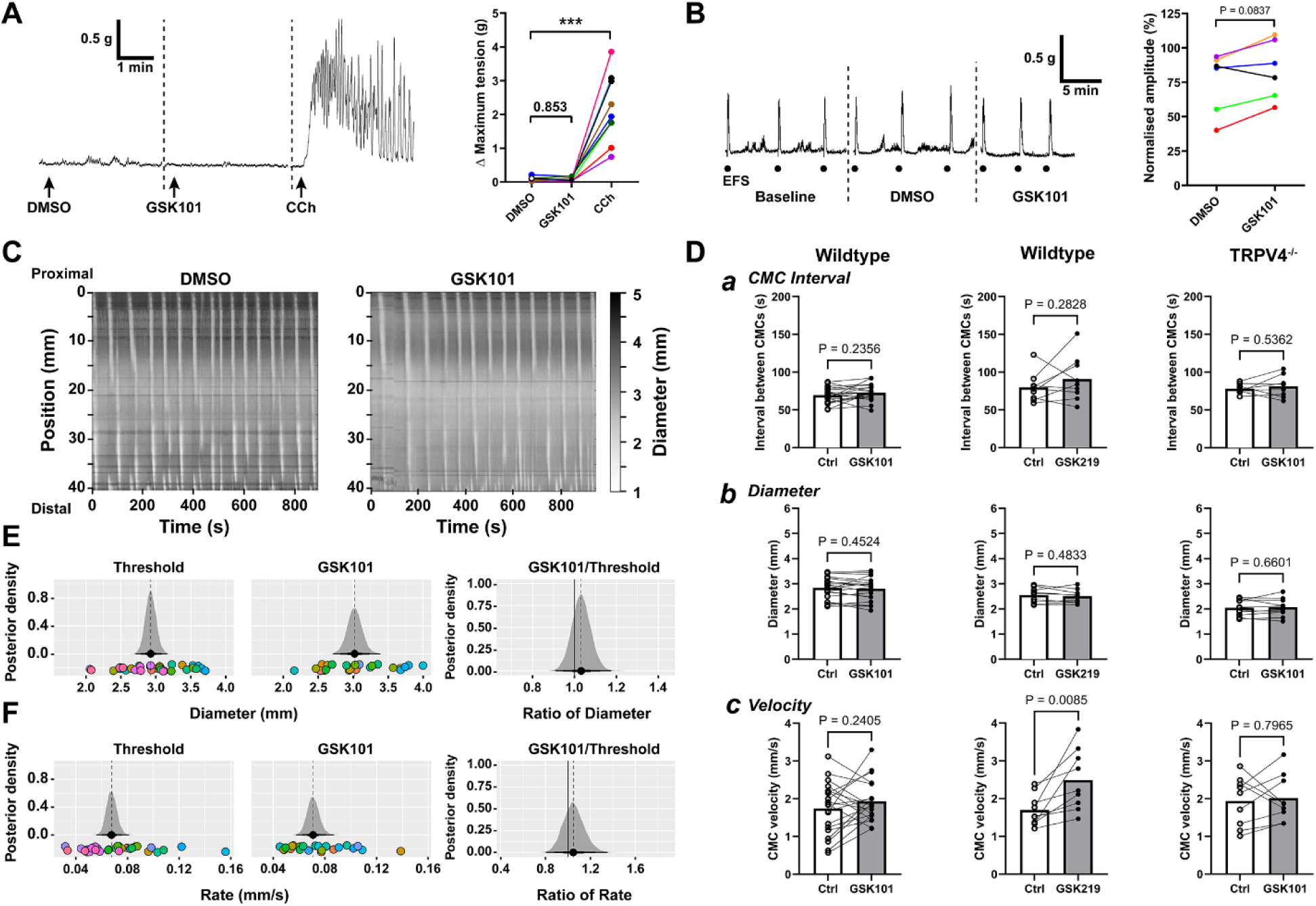
TRPV4 activation does not affect contractility of the isolated mouse colon. **A.** Representative trace demonstrating no effect of GSK101 (100 nM) on circular muscle tension relative to vehicle (DMSO). Tissues contracted in response to the positive control carbachol (CCh, 10 µM; ***p<0.001, n=9 mice, one-way ANOVA, Tukey’s post-hoc test). **B.** TRPV4 activation (GSK101, 100 nM) had no effect on the mean amplitude of electrically evoked neurogenic contractions relative to DMSO control. A representative trace of responses to EFS (•) recorded at baseline and following subsequent addition of DMSO and GSK101 (100 nM) is shown (p=0.084, n=6, paired t-test). **C.** Representative DMaps showing propagating contractions (vertical white lines) under threshold intraluminal pressure in the presence of vehicle (DMSO, left) then GSK101 (100 nM, right). **D. (*a*)** The interval between CMCs and mean colon diameter **(*b*)** was unaffected by pharmacological activation (*left*: 100nM GSK101) or inhibition (*middle*: 1µM GSK219) of TRPV4, or by TRPV4 knockout (*right*). **(*c*)** CMC velocity was not changed in response to GSK101 (*left*) or TRPV4 deletion (*right*) but was significantly enhanced in the presence of GSK219 (*middle*). Points represent mean values per preparation, n=9-20 mice per group. Data were compared by paired (GSK101, GSK219) or unpaired (TRPV4^-/-^) two-tailed t-tests. **E.** Posterior distribution graphs demonstrate the probability of a contraction occurring at different colon diameters under control or GSK101-treated conditions. Colored dots represent data from independent experiments (n=34 and 24 mice). No difference in mean gut diameter was detected between control and GSK101-treated preparations (ratio of the mean diameter= 1.03). **F.** Posterior distribution graphs summarizing the mean rate of contraction under control and GSK101-treated conditions (n=34 and 24 mice). There was no difference in the rate of contractions between the two groups (ratio of the mean= 1.04).

Experimental and clinical intestinal inflammation is consistently associated with upregulation and sensitization of TRP channels, including TRPV4 [10, 17]. Myenteric neurons of the acutely inflamed mouse colon did not respond directly to GSK101 (n=369/369 neurons, N=7 mice), suggesting that unmasking of latent TRPV4 activity through upregulation and sensitization of the channel [8] does not occur in enteric neurons under these pathophysiological conditions.

### TRPV4 has minimal influence on motility of the mouse colon

TRPV4 activation may promote either contraction or relaxation of the mouse colon [17, 19]. To clarify the role of TRPV4 in motility, we first measured contractile activity of isolated colon segments treated with GSK101. Vehicle (0.01% DMSO) or GSK101 (100 nM) did not alter smooth muscle tension within 2 min of bath application (vehicle: 0.08 ± 0.06 g vs GSK101: 0.08 ± 0.06 g, n=9 per group, mean ± SD, **Fig. 6A**). GSK101 had no effect on phasic activity or tone, consistent with the lack of prominent Ca^2+^ responses to TRPV4 activation by myenteric neurons, ICC or PDGFRα cells. All tissues contracted in response to CCh (10 µM), which served as a positive control for tissue viability. The amplitude of electrically evoked neurogenic contractions was unaffected by GSK101 treatment (vehicle: 82.2 ± 27.1%, GSK101: 94 ± 32.6%, n=7; **Fig. 6B**). The lack of effect of TRPV4 modulation on neurogenic contractions is consistent with the inability to detect functional expression of TRPV4 by myenteric neurons using Ca^2+^ imaging.

### Lack of prominent effect of TRPV4 in propagating colonic motility

The influence of TRPV4 activity on propagating motility patterns was assessed using the intact isolated mouse colon. Application of GSK101 (100 nM) to selectively activate TRPV4 did not affect the time interval between CMCs or CMC velocity (n=20, **Fig. 6C & D**). The resting colon widths between CMCs were compared as a reflection of baseline muscle tone. GSK101 had no effect on the resting width of the proximal or distal regions of the colon (n=20, **Fig. 6D**). Inhibition of TRPV4 (GSK219, 1 µM) significantly increased CMC velocity, but had no effect on the time interval between CMCs (n=9, paired data, **Fig. 6D**). The resting gut width between contractions was also unaffected, consistent with a lack of TRPV4 involvement in setting basal tone (N=11 TRPV4^-/-^, N=20 WT). TRPV4 deletion (TRPV4^-/-^ mice) was not associated with a significant change in CMC interval or velocity when compared to wildtype (WT) colons (N=9 TRPV4^-/-^, N=9 WT). However, the resting widths of TRPV4^-/-^ colons were significantly narrower than WT controls (one-way ANOVA). The interval between CMCs, CMC velocity, and colon diameter were not significantly different in TRPV4^-/-^ colons following GSK101 addition (n=8) (**Fig. 6D**). These data do not support a prominent role for TRPV4 in the physiological control of motility.

### TRPV4 modulates distension-evoked colonic contractions

The potential role of TRPV4 as a mechanosensitive ion channel that responds to distension of the gut wall was assessed using spatiotemporal mapping by measuring the *mean diameter of contractions* (MDC), defined as the mean colon diameter immediately prior to initiation of a CMC. The gut wall was distended by increasing intraluminal pressure and responses were compared with those recorded from the same preparation at baseline pressure. Increasing intraluminal pressure increased the MDC (baseline diameter: 2.29-2.64 mm vs. threshold diameter: 2.60-2.97 mm, **Fig. S2**), confirming that differences could be detected using this analysis method. Bath addition of GSK101 (100 nM) at threshold had no effect on the MDC (vehicle: 2.78-3.08 mm vs. GSK101: 2.81-3.24 mm, ratio of mean 1.03, **Fig. 6E**). The MDC in the presence of GSK219 ranged from 2.35-2.82 mm, with no difference in the ratio of the mean diameter in the presence of GSK219 (0.93) (**Fig. S14A**).

To assess how TRPV4 influences the sensitivity of the colon to distension we used TRPV4^-/-^ mice and pharmacological modulators of channel activity. The MDC of TRPV4^-/-^ or WT colons were compared at baseline and threshold intraluminal pressures. The MDC at baseline pressure was lower in TRPV4^-/-^ colons compared to WT controls (TRPV4^-/-^: 1.8-2.1 mm vs. WT: 2.3-2.6 mm, ratio of means= 0.78, **Fig. S15A, B**). Similarly, the MDC at threshold pressure was smaller in TRPV4^-/-^ colons compared to WT (TRPV4^-/-^: 2.16-2.70 mm vs. WT 2.61-2.96 mm, ratio of means= 0.87). At threshold, the MDC in the presence of GSK101 was lower in tissues from TRPV4^-/-^ mice compared to WT mice (ratio of means=0.82). This may reflect a difference in the responsiveness of TRPV4^-/-^ tissue to threshold stimulation, as GSK101 treatment did not have any effect when compared to vehicle control (ratio of means=0.97). The MDC for TRPV4^-/-^ colons was greater at threshold than at baseline (ratio of means=1.30) (**Fig. S15C**).

The role of TRPV4 in influencing the onset of contractions was examined by measuring the *mean rate of contractions* (MRC; *see Methods*). Under threshold conditions there was no difference in the MRC following GSK101 treatment when compared to vehicle control (vehicle: 0.06-0.08 mm/s vs. GSK101: 0.06-0.08 mm/s, ratio of rate: 1.05, **Fig. 6F**). However, the MRC was lower in the presence of GSK219 when compared to vehicle (vehicle: 0.05-0.07 vs. GSK219: 0.04-0.05, ratio of rate =0.77) (**Fig. S14B**). At baseline intraluminal pressure, the MRC was lower in tissue from TRPV4^-/-^ mice compared to WT colon (TRPV4^-/-^: 0.03-0.04 mm/s vs WT: 0.05-0.06mm/s, ratio of means=0.67, **Fig. S16A, B**). No differences in the MRC between WT and TRPV4^-/-^ tissues was detected at threshold (ratio of rate=0.90) (**Fig. S16C**).

Collectively, these data demonstrate that TRPV4 plays a role in activation of contractile responses to mechanical distension of the colon. However, TRPV4 has minimal involvement in the active initiation or regulation of propagating colonic motor patterns.

## Discussion

TRPV4 is an emerging therapeutic target for inflammatory disease and pain [8, 9]. Although TRPV4 inhibitors have potential for the treatment of IBS and IBD symptoms [47], fundamental knowledge of the functional roles of TRPV4 in different cell types in the GI tract is either lacking or contradictory. This includes consensus of where TRPV4 is expressed. The present study identified unique populations of cells responsive to TRPV4 activation in the colon and provided greater mechanistic insight into the involvement of TRPV4 in GI functions including motility. We present evidence that TRPV4 activation evokes pronounced Ca^2+^ transients in enteric glial cells, macrophages, and lymphatic and vascular endothelia. This information has implications for how TRPV4 may contribute to intestinal inflammation and reveals possible roles for TRPV4 in interstitial fluid homeostasis, effective immune presentation to draining lymph nodes, and for localized production and removal of inflammatory mediators.

### TRPV4 does not play a prominent role in the neurogenic control of colonic motility

A key finding of this study was the lack of prominent responses to TRPV4 activation by enteric neurons. Although previous studies have reported that TRPV4 is functionally expressed by myenteric and submucosal neurons of the intestine [17, 18, 20], we found no clear evidence to support functional expression of TRPV4 by enteric neurons in the mouse, monkey, or human colon using a range of complementary approaches. TRPV4-dependent activation of neurons was not detected within the resolution of our recording apparatus in tissues using either calcium-sensitive dyes or genetically encoded biosensors. Similarly, no activation of myenteric neurons by GSK101 was observed under culture conditions, where phenotypic changes may occur. GSK101-evoked Ca^2+^ responses by other cell types persisted in the presence of TTX and were replicated using isolated cells, consistent with direct activation. Furthermore, there was no evidence for neuronal activation in acutely inflamed tissues, suggesting that unlike in nociceptive pathways [8, 10], pathophysiological upregulation or sensitization of TRPV4 in enteric neurons is unlikely to occur. This differs to evidence that prior exposure to the proinflammatory mediator histamine enhanced TRPV4 signaling in human submucosal neurons through the H1 histamine receptor [18]. Our observations are consistent with single cell transcriptomic data which indicate limited expression of TRPV4 mRNA in neurons of the mouse colon [48] and with Matheis *et al*. (2020) [49] who used SNAP25Cre-RiboTag mice to sequence translating RNA from myenteric neurons. Furthermore, we demonstrated that TRPV4 activation, pharmacological block, or genetic deletion did not affect neuromuscular function of the colon or its capacity to generate and coordinate propagating motor patterns. This is consistent with the inability to detect TRPV4-dependent activation of enteric neurons and smooth muscle cells. TRPV4 activity did not influence phasic activity or tone, and Ca^2+^ imaging studies indicated that TRPV4 was not functionally expressed by ICC or PDGFRα cells. These observations do not align with evidence that activation of TRPV4 can inhibit [17] or enhance [19] GI motility. Our studies have identified a novel role for TRPV4 as a detector of elevated intraluminal pressure or tissue distension, and as an initiator of coordinated propagating motility. TRPV4 appears to set the threshold at which CMCs are evoked but is not essential for this process, as evidenced by the retention of CMCs in TRPV4^-/-^ tissues and following treatment with the highly potent TRPV4 antagonist GSK219. Although the precise cellular mechanisms involved remain to be determined, they may involve activation of TRPV4 expressed by mucosal cell types not specifically examined in this study, glia, or extrinsic afferents.

### The Type I enteric glial population is heterogeneous

The current classification scheme on which glial populations are defined is based on their location and organization within the gut wall. However, recent findings including those presented in the current study provide compelling evidence that the glial populations within each subtype are highly heterogeneous [48, 50, 51, 52]. This is consistent with recent evidence for heterogeneity in satellite glial populations in sensory and sympathetic ganglia [53]. Our connectivity analysis has identified a subpopulation of Type I glia that may reflect a TRPV4-positive ‘initiator’ population that promotes sustained signaling within the glial network. This conclusion is based on the kinetics of responses to GSK101 and by the effects of gap junction inhibition. We propose an analogous model to that identified in astrocyte populations, in which TRPV4-dependent activation of a subset of cells is sufficient to drive intercellular coupling and reinforce sustained activity within the network [38]. However, the mediators involved appear to differ between the two cell types [38, 54]. Our model is consistent with the effect of gap junction inhibition on the duration of TRPV4-dependent signaling by a subpopulation of glia. PCF analysis of the effects of TTX revealed that neuronal input is required for the reinforcement of glial network activity under the conditions examined.

TRPV4 promoted ATP release from astrocytes and block of purinergic signaling abolished residual oscillatory activity in the presence of gap junction inhibitors [38]. In contrast, block of purinoceptors dampened coordinated activity but did not alter temporal Ca^2+^ signaling in the enteric glial network. Ca^2+^ influx following TRPV4 activation can lead to Ca^2+^-induced Ca^2+^ release in astrocytic endfeet [39]. Depletion of intracellular Ca^2+^ stores in enteric glia using the SERCA inhibitor CPA had no effect on the average number of peaks but resulted in a significant increase in rise-time in an analogous manner to the slow kinetics of TRPV4 activity in CPA-treated astrocytic endfeet [39]. However, this does not explain why CPA pre-treatment resulted in a significantly higher proportion of glia that responded to GSK101. The inhibition of Cx43 hemichannels disrupted the TRPV4-induced coordinated activity of the glial network. It is possible that TRPV4-activation led to the release of ATP via Cx43 hemichannels, resulting in paracrine activation of glia and propagation of Ca^2+^ waves. This is supported by the disruption of glial signaling by purinoceptor block as revealed by PCF analysis, and by similar studies in astrocytes demonstrating ATP-release via Cx43 hemichannels after TRPV4 activation [38, 54]. Sustained oscillatory activation of the glial network in response to the neuropeptide neurokinin A is also suppressed by TTX and glia-specific conditional deletion of Cx43 hemichannels [40], in line with the observations that we report.

### Limitations of the current study

Endogenous activators that are likely to contribute to TRPV4 signaling include lipid derivatives and local changes in tonicity. Although our functional distribution studies have relied exclusively on the use of the small molecule TRPV4 activator GSK101, this was an essential first step for validating our approach and key findings. GSK101 is a highly potent and selective activator of TRPV4, as confirmed by the absence of Ca^2+^ responses in TRPV4^-/-^ tissues and in the presence of TRPV4 inhibitors. GSK101 was selected for its ability to robustly activate TRPV4 in all cells with established TRPV4 expression. This approach reduced the potential for off-target effects and the associated complexity of applying non-selective agonists or unfocused activators such as endogenous lipids or mechanical stimuli. It is also noted that this study focuses exclusively on TRPV4-mediated Ca^2+^ signaling and does not assess other downstream signaling pathways, the longer-term effects of sustained TRPV4 activation on the colon, or TRPV4-dependent mediator release.

This study has broader implications for the involvement of other cell types in colonic motility. GSK101 treatment resulted in prolonged elevations in intracellular Ca^2+^ in all mMac and sustained Ca^2+^ oscillations in the glial network. These prominent responses were not reflected in the effects of GSK101 on contractile activity or coordinated motility, suggesting that mMac and glia have minimal influence on motility under the assay conditions examined. These observations contrast with recent evidence highlighting the importance of both mMac and glia for normal colonic motility [19, 34, 45, 55]. The use of mice with a global and non-inducible deletion of TRPV4 partly limits the depth of mechanistic insight provided by motility studies. For example, it is possible that deletion of TRPV4 from cells with opposing roles may result in a lack of net effect on motility. A similar consideration must be made when interpreting motility studies using GSK101, which robustly activates all TRPV4 expressing cells in the colon.

TRPV4 also influences the magnitude and duration of GPCR and ion channel signaling. TRPV4 can be sensitized by phosphorylation and activated by lipid intermediates downstream of GPCR activation [8, 56]. Although this is a major role for TRPV4, particularly in pathophysiology, the involvement of TRPV4 as a modulator of signaling by these receptors was not examined in the current study. However, the detailed information of where TRPV4 is expressed provided by this study combined with existing information about the GPCRs that are co-expressed opens new avenues for future exploration.

In summary, our observations reveal new insight into how TRPV4 may influence gut function in health and disease. We provide new approaches to assess TRPV4 signaling and new information about where TRPV4 is expressed. Our consistent findings across species indicate potentially conserved functions for TRPV4 in the colon. We conclude that TRPV4 is not a primary mechanosensory in enteric reflex pathways that coordinate GI motility.

## Acknowledgements

Taro Ishikawa, (Jikei University School of Medicine: TaroTools)

Stephen Kent, Isaac Barber-Axthelm (The Peter Doherty Institute for Infection and Immunity: macaque tissue)

Jakub Fichna (Medical University of Lodz: initial studies of TRPV4 in the gut) Wolfgang Liedtke (Duke University: trpv4-/- mice)

Lee Travis (Flinders University: assistance with Ca2+ imaging)

Heather Young, Lincon Stamp, Marlene Hao (The University of Melbourne: wnt1-GCaMP3 mouse tissue)

## Funding

National Health and Medical Research Council Australia (NHMRC) 1083480 (DPP), Australian Research Council (ARC) Centre of Excellence in Convergent Bio-Nano Science and Technology (CE140100036, EJC, NAV), ARC Discovery Project (DP170101358, EJC), ARC Future Fellowship (FT220100617, NAV), ARC Discovery Early Career Researcher Award (DE200100988, STJ; DE200100825, SEC), NIDDK grants R01DK103723 and R01DK120862 (BDG)

Research in DPP, NAV and SEC’s laboratory was funded in part by Takeda Pharmaceuticals.

## SUPPLEMENTAL INFORMATION

### Supplementary Methods Immunostaining post calcium imaging

Wholemounts were fixed immediately after Ca^2+^ imaging (4% paraformaldehyde (PFA), overnight, 4°C). They were then washed (3x PBS), blocked (blocking buffer: 5% normal donkey serum in PBS with 0.1% Triton X-100) (1h, RT; human tissue: overnight, 4°C), then labeled with primary antibodies (in blocking buffer, 2-3 days, 4°C; human tissue: 5 d; **Table S3**). Preparations were washed with PBS (3x) and incubated with fluorophore-conjugated secondary antibodies (1:500 in PBS, 1 h, RT; human tissue: overnight, 4°C; ThermoFisher). Tissues were washed (3x PBS), labeled with the nuclear marker DAPI (1:1000 in PBS, 5 min; human tissue: 1:500, 2 h), then mounted using ProLong Diamond Antifade mountant (ThermoFisher) or buffered glycerol (human tissue). Images were captured with a Leica TCS SP8 laser-scanning confocal microscope (20x HC PLAN APO NA 0.75) or a Leica TCS SP8 Lightning confocal system (20x HC PL APO NA 0.88 or 40x HC PL APO NA 1.1 water immersion objective). Images were acquired at 4096 x 4096 or 2048 x 2048 resolution at 16-bit depth.

### Image processing and analysis

All images were processed using FIJI (ImageJ v1.52c) [1, 2]. “*Linear stack alignment with SIFT*” [3] or the “*Template Matching and Slice Alignment*” plugin were used to register image stacks from Ca^2+^ imaging. The immunolabeled images were overlaid on the maximum projection of Ca^2+^ imaging frames, and non-linear tissue deformation due to the immunostaining process was accounted for using the “*bUnwarpJ*” plugin [4]. The overlaid immunolabeled image enabled specific identification of distinct cell types and was used as a mask for generating regions of interest (ROIs) (e.g., Sox10+ enteric glial cells). ROIs were also identified based on the Ca^2+^ response profiles of cells to pharmacological activators (e.g., MRS2365 and TFLLR-NH_2_ responsive PDGFRα+ cells). In initial experiments, the identity of lymphatic vessels, blood vessels, and macrophages was confirmed by immunostaining for LYVE1, CD31 and mMac specific markers, respectively. Cells were readily identifiable in subsequent experiments based on morphology and their Ca^2+^-responses to GSK101. TRPV4-mediated Ca^2+^ responses by endothelia and macrophages were rapid in onset and sustained, allowing their use as internal controls to indicate effective agonist exposure. This combination of approaches enabled the accurate selection of ROIs and analysis of response profiles by specific cell populations. Cells were excluded if they were outside of the focal plane or located at the edge of the image.

Image series were normalized to baseline (F/F_0_), which was defined as the average of 30 frames prior to GSK101 addition. The mean intensity values for every timepoint were extracted for each ROI and used for quantitative analysis.

### Data processing and analysis

Datasets were imported into Igor Pro 8.0.2.1 (Wavemetrics, Portland, OR, USA) for analysis of Ca^2+^ responses. To extract features from these waves, a set of procedure files (“TaroTools”: https://sites.google.com/site/tarotoolsregister/) were used. The peak Ca^2+^ responses were defined as ‘*events*’ and each cell as an ‘*episode*’ within TaroTools. The “*Event Detection*” window was used with the following parameters:

- Threshold: 0.1
- Smooth1: 3
- Baseline was selected as “*Polarity-dependent horizontal*”.
- Basepoints: 0
- Time frame for analysis was set to 240 s post drug addition.

False events due to focal drift were selected using “*Mask event*” and deleted using “*Delete Masked*”. “*Conditional Mask*” was used to mask false events that were present across all traces following drug addition or tissue movement. “*Check Errors*” verified any errors in the detection of parameters. The “*Further analysis*” option was applied to extract the following features from each Ca^2+^ wave trace:

- Events (number of Ca^2+^ peaks per cell)
- Event times
- Event onset times
- Amplitude
- Area under the curve (AUC)

### Temporal color coding

The “*Temporal color coding*” plugin (Image-> Hyperstacks) in FIJI was used to visualize the timing of Ca^2+^ responses. To ensure that responsive cells were brighter and to reduce noise, the macro (“*Temporal-Color_Code.ijm”*) bundled with FIJI was modified to include “*Standard Deviation*” as an option for the Z-projection method.

### Pair correlation analysis

Pair correlation functions (PCFs) were calculated using the coordinates of the EGCs, their Ca^2+^ peaks and the area encompassing the ganglia. EGC coordinates were taken from the centroid of each cell and ROIs were extracted within FIJI. Outlines around ganglia were manually drawn for each image and the coordinates extracted to define the ganglionic borders. The distance and adjacency matrix were used to calculate the PCFs [5, 6]. The PCF was first calculated from EGCs that were activated within 30 s after treatment application and then this process was repeated for every subsequent 30 s period. If the PCF value was greater than 1 at distance *x* at time *t* then there were more pairs of responding EGCs separated by distance *x* at that time than expected if responses were random (i.e., EGC activity is coordinated). Similarly, if the PCF value was less than 1 at distance *x* then there were fewer pairs of activated glial cells separated by that distance than expected if cells were activated randomly. PCF values above and below 1 at *x* correspond to positive and negative correlation, respectively, between pairs of cells separated by a distance *x* (**Fig. S1D, E**).

### Primary cell culture and cell lines

#### RAW264.7 murine macrophage cell line

RAW264.7 cells (ATCC: TIB-71) were maintained in Dulbecco’s Modified Eagle Medium (DMEM) containing 10% FBS and penicillin/streptomycin according to ATCC guidelines. Cells were seeded onto 96-well plates (20,000 cells/well) and cultured for 24h before use in population Ca^2+^ assays.

#### Enteric glial cell line

The rat enteric glial cell line EGC/PK060399egfr: (ATCC CRL-2690) [7] was maintained in DMEM containing 10% FBS and penicillin/streptomycin with 1% G-5 supplement (ThermoFisher). Cells were seeded onto 96 well plates (14,000 cells/well) coated with poly-D-lysine (100 μg/ml) and cultured for 24h before use in population Ca^2+^ assays.

#### Primary human lymphatic endothelial cells

Human adult dermal lymphatic endothelial cells (HMVEC-dLyAd-Der Lym Endo Cells; Lonza) were maintained using EBM-2 Basal Medium supplemented with EGM-2 MV Microvascular Endothelial Cell Growth Medium supplements (Lonza). Cells were maintained for 8 or fewer passages in a T75 flask. Cells were seeded onto 96 well plates at 30,000 cells/well and cultured for 24 h before use in population Ca^2+^ assays.

#### Isolation of peripheral blood mononuclear cells (PBMCs) and differentiation into human monocyte derived macrophages

Blood collected from 3 healthy blood donors by the Australian Red Cross was used for PBMC preparation followed by monocyte isolation as described previously [8]. Briefly, whole blood (100 mL) was collected into K2EDTA vacutainer tubes (BD). Whole blood was diluted with PBS (1:1; 30 mL total volume) and overlaid onto 15 mL Lymphoprep (Stem Cell Technology) at RT. Blood was centrifuged at 160 *g* for 20 min with the centrifuge brake set to low, followed by removal of 20 mL of supernatant then centrifugation of the remainder at 350 *g* for 20 min. PBMCs were collected from the interphase and washed twice by centrifugation in PBS (with 2% FBS, 1 mM EDTA) at 150 *g* for 15 min. Red blood cells were lysed using ACK lysis buffer (Gibco) for 3 min, then remaining cells were washed with PBS.

Monocytes were isolated from PBMCs using isosmotic Percoll (9 parts Percoll to 1 part 1.5 M NaCl) mixed in equal proportions with PBS/citrate (NaH_2_PO_4_ 1.49 mM; Na_2_HPO_4_ 9.15 mM; NaCl 139.97 mM; C_6_H_5_Na_3_O_7_ .2H_2_O 13 mM; pH 7.2) [9]. PBMCs were resuspended in PBS at a density of 1-2x10^7^ cells/ml and 5 mL of this cell suspension was gently overlaid onto 9 mL of Percoll/PBS/citrate solution and centrifuged at 400 *g* for 35 min with low brake. The monocytes were collected from the interface and washed twice with PBS prior to resuspension in RPMI-1640 culture medium (2 mM L-glutamine, 100 U/ml penicillin and 100 μg/ml streptomycin) supplemented with 10% FBS. The monocytes were cultured in the presence of 5000 U/mL of human M-CSF for 7 d to differentiate them into macrophages. On day 3, the non-adherent cells were aspirated, and adherent cells replated onto 96 well plates at a density of 30,000 cells/well in RPMI-1640 media with 10% FBS and M-CSF. Monocyte-derived macrophages were cultured for another 4 d before use in population Ca^2+^ assays.

#### Mouse bone marrow-derived macrophages

Mouse bone marrow-derived macrophages were cultured as previously described [10]. GM-CSF bone marrow-derived macrophages (GMBMDMs) were cultured in the presence of murine GM-CSF (20 ng/mL). M-CSF bone marrow-derived macrophages (BMDMs) were cultured with murine M-CSF (5000 U/mL). Cells were seeded onto 96 well plates (50,000 cells/well) coated with poly-D-lysine (100 μg/ml) and cultured for 24 h before use in population Ca^2+^ assays.

#### Primary culture of myenteric neurons of the mouse colon

The mouse colon was removed and placed in cold HBSS containing antibiotic/antimycotic solution (1% v/v; 10,000 units penicillin, 10 mg streptomycin and 25 μg amphotericin B per mL; Sigma-Aldrich) and nicardipine (1 µM)). The colon was opened along the mesenteric border and flushed of contents before stretching and pinning mucosa downwards onto a silicone elastomer-lined dish. The external muscle was isolated by sharp dissection, then incubated in Ca^2+^-Mg^2+^-free Hanks’ balanced salt solution containing 1 mg/mL collagenase II and IV and 0.1 mg/mL DNaseI for 45 min. Tissue was dissociated by mechanical trituration using fire-polished Pasteur pipettes. The resulting cell suspension was washed by repeated centrifugation (2 x 500 *g,* 5 min, resuspension in DMEM). Dissociated cells were plated onto wells of a μ-Slide 8 Well Grid-500 chamber slide (ibidi GmbH, Martinsried, Germany) coated with poly-L-lysine and 100 μg/mL laminin (Sigma-Aldrich). Cells were maintained in DMEM containing antibiotic-antimycotic, 10% FBS and 1% N-1 neuronal supplement (Sigma-Aldrich) in a humidified incubator at 37°C (95% O_2_, 5% CO_2_) for 7 d to allow enteric glial cells to proliferate. For primary myenteric neuron cultures, cytosine arabinoside (1 µM; Sigma-Aldrich) was added to the complete culture medium to prevent the growth of non-neuronal cells. Neurons were studied after 5-7 d *in vitro* [11].

### Measurement of intracellular Ca^2+^ signaling

#### Plate-based Ca^2+^ assays

Cells were loaded with Fura-2 AM (3 μM, 45 min, 37°C) in Ca^2+^ assay buffer (10 mM HEPES, 0.5% BSA, 10 mM D-glucose, 2.2 mM CaCl_2_.H_2_O, MgCl_2_.6H_2_O, 2.6 mM KCl, 150 mM NaCl) containing 4 mM probenecid and 0.05% pluronic F127. Fluorescence was measured at 4 s intervals (excitation: 340 nm/380 nm; emission: 520 nm) using a FlexStation 3 plate reader (Molecular Devices, Sunnyvale, CA). A baseline recording (16 s) was made before cells were challenged with GSK101 (500 pM to 10 μM). In some experiments, cells were pre-treated for 30 min with 10 µM HC067047 to assess the TRPV4-dependence of responses. Ionomycin (1 μM) was applied at the end of each experiment as a positive control [12].

#### Ca^2+^ imaging and characterization of primary cells

Cells were loaded with Fura-2 AM as described above. Cells were washed and incubated in Ca^2+^ buffer for 20 min before imaging. A Leica DMI-6000B microscope with an HC PLAN APO 0.4 NA x10 objective and incubator system (37°C) was used for Ca^2+^ imaging. The cells were challenged with 100 nM GSK101 and 10 µM ATP and images were collected at 1 s intervals (excitation: 340 nm/380 nm; emission: 530 nm) [12]. Cells were fixed (4% PFA, 4°C, 10 min) immediately after Ca^2+^ imaging. They were then washed (3x PBS), blocked (5% normal horse serum in PBS with 0.1% saponin, 1h, RT), then incubated with primary antibodies (in blocking buffer, overnight 4°C). Cells were subsequently washed (3x PBS), incubated with secondary antibodies (1 h, RT) and DAPI (5 min in PBS), then washed (3x PBS). Images were captured with a Leica TCS SP8 laser-scanning confocal microscope using an HC PLAN APO 1.3 NA x40 or 1.4 NA x63 oil immersion objective. In some experiments, macrophages were labeled directly with anti-F4/80-Alexa488 prior to Ca^2+^ imaging.

### Macrophage depletion

Muscularis macrophages were depleted using CSF1R blocking antibodies. Briefly, mice were treated with either anti-CD115 or rat IgG2A (18.75 µg/g, i.p. in final volume of 0.2mL sterile 0.9% saline, days 0 and 1; [13]). Mice were killed on day 3 and wholemounts of the distal colon were prepared for Ca^2+^ imaging. Effective depletion or retention of mMac was confirmed for all mice by immunostaining with combinations of antibodies (CD68, CD169, MHCII, and IBA1) to account for potential loss of marker expression, rather than cell loss. The presence of mMac in mAb treated tissues was used as grounds for exclusion from analysis.

### TNBS-induced colitis model

Acute colitis was induced in C57Bl/6 mice according to methods outlined in Wirtz *et al*. 2017 [14]. Mice were pre-sensitized by application of 1% TNBS (5% picrylsulfonic acid solution (Sigma-Aldrich); w/v in 4:1 acetone: olive oil vehicle) to the nape skin (1 part 5% TNBS stock in 4 parts vehicle). After 8 days, TNBS (2.5% picrylsulfonic acid solution in 1:1 ethanol and 0.9% saline) was administered to the colon. Mice exhibited weight loss, loose stools, and rectal bleeding consistent with colitis. The mean weight loss on d3 post TNBS was 8.08 ± 1.36% (n=8 mice). Mice were humanely killed by cervical dislocation 3 days after TNBS induction. Inflammation was confirmed by macroscopic damage, and by increased infiltration of CD45 and CD68 cells into the mucosa, and by increased density of MPO (rabbit anti-myeloperoxidase, A0398, DAKO, RRID:AB_2335676) and 3NT (rabbit anti-nitrotyrosine, 06-284 Sigma-Aldrich, RRID:AB_310089) labeling.

### Tissue contraction assays with electrical field stimulation (EFS)

Mouse colons were collected in modified Krebs buffer (in mM: 118 NaCl, 4.7 KCl, 7 MgSO_4_, 1.1 H_2_O, 1.18 KH_2_PO4, 25 NaHCO_3_, 11.6 glucose, and 2.5 CaCl_2_). The proximal colon was cut into segments (5 mm lengths), mounted onto holders embedded with stimulating electrodes, then placed under 0.5-1 g tension in 10mL water-jacketed organ baths (Krebs buffer, 37°C, bubbled with 95% O_2_/ 5% CO_2_, [15, 16]). Circular muscle tension was detected by a force transducer (FT03, Grass Instruments, Quincy, MA, USA). Recordings were captured with a PowerLab 4/SP system (ADInstruments, Australia) and viewed using LabChart software (Version 5.0.2). Stimulating electrodes were connected to Grass S88 stimulators (Grass Instruments, MA, USA) and transmural electrical field stimulation was used to evoke neurogenic contractions (EFS; 0.5 ms duration, 60 V, 1000 ms train duration, 3 pulses s^-1^; 5 min interval between stimulations). Baseline activity was defined once contractions to EFS were reproducible (≥30 min post dissection). Tissues that generated <0.2 g tension above baseline were excluded from the analysis. Carbamoylcholine (carbachol, CCh; 10µM) was added at the conclusion of each experiment to confirm tissue viability. Unresponsive preparations were excluded from analysis.

Tissues were treated with vehicle (0.001% DMSO) and then GSK101 (100 nM). The amplitude of contractions within 2 min after drug addition was compared to the baseline to determine whether the addition of GSK101 stimulated contractile activity. The mean amplitude of three EFS-evoked contractions (5 min post-treatment) was compared for each condition. The average amplitude of EFS-induced contractions was normalized to baseline responses to determine the effect of TRPV4 modulation on neurogenic contractions.

### Colonic motility assays

Video recordings of diameter changes were used to analyze motor patterns generated by the isolated mouse colon [17]. The excised colon was placed into a customized, water-jacketed chamber continuously perfused with Krebs buffer (37°C). The colon was cannulated at both ends via cotton ligatures. An inflow reservoir containing 20mL of Krebs buffer was connected to the proximal cannula and set to the height of the colon (baseline pressure). Back-pressure was maintained via an outflow tube connected to the distal cannula and the height of fluid within this outlet was maintained at 2cmH_2_O. Intraluminal pressure was increased by raising the height of the inflow reservoir to a point where consistent propagating contractions could be observed (threshold, 10-12 cm). A camera positioned above the colon captured motor patterns during the experiment (Logitech QuickCam Pro; 15 FPS, VirtualDub software Version 1.10.4).

Tissues were equilibrated for 30 min at baseline intraluminal pressure. Colonic activity was recorded at baseline for 10 min, then threshold pressure for 10-15 min, followed by 10-15 min at threshold in the presence of either GSK101 (100 nM) or GSK219 (1 µM). After washout (5 min), CCh (10 μM) was applied directly to the bath to confirm tissue viability.

#### Analysis of colonic migrating complexes (CMCs)

To allow the analysis of changes in diameter, video files (.AVI) were converted into diameter maps (DMaps) via an in-house edge detection software (MATLAB software, version R2019a Update 5, [18]). Specific CMC parameters were analyzed using custom tools built into this software. CMCs were defined as a contraction that travelled more than 40% of the total colon length. As an indicator of the frequency of contractions, the time interval between the start of each CMC was calculated and averaged for the duration of the test condition. The resting diameter of the colon was determined by averaging the diameter at randomly selected points between contractions. The resting diameter was calculated across the proximal and distal regions of the colon (25% and 75% of the colon length). Each spatiotemporal location on the DMap was labelled as either ‘contracting’ or ‘non-contracting’ based on whether the negative rate of change of diameter with respect to time surpassed a constant threshold. A graphical overview of the analysis used is presented in **Figure S2**.

#### Statistical analysis of CMC data

For standard parameters, data were analyzed using Prism 8 (GraphPad Software, San Diego, CA). The results were expressed as mean ± standard deviation (SD). The normal distribution of the data was determined through the Shapiro-Wilk test or Kolmogorov-Smirnov test. Any outliers within the data were identified through the ROUT method in Prism 7 (Q=1%) and excluded from further analysis. The data were analyzed by two-tailed Student’s t-test or one-way ANOVA. A p value of less than 0.05 was defined as statistically different to the null hypothesis at the 95% confidence level. The brms R-package [19] was used to analyze changes in diameter and the rate of contraction (**Figure S2**). There were two categorical variables: *cond* refers to the environmental condition of the preparation, and *type* refers to the nature of the gut.

The conditions variable could be in one of the four levels: *Baseline*, *Threshold*, *GSK101*, and *GSK219*. GSK101 or GSK219 refer to the periods at threshold pressure in the presence of either GSK101 or GSK219 respectively. The formula for the mu parameter was given by “mvbind(log(diam), log(rate)) ∼ cond * type”, with sigma parameter given by “cond * type”, under a Gaussian family with residual correlations. Default priors were used for intercepts, and a “normal(0,1)” was applied prior to the “b” coefficients. 8-chains of 3000 iterations were used, which included 1000 warm-up iterations, and an adapt-delta of 0.99. No problems with the fit were detected with 0 divergences and an Rhat very close to 1.

---

**Table S1.**
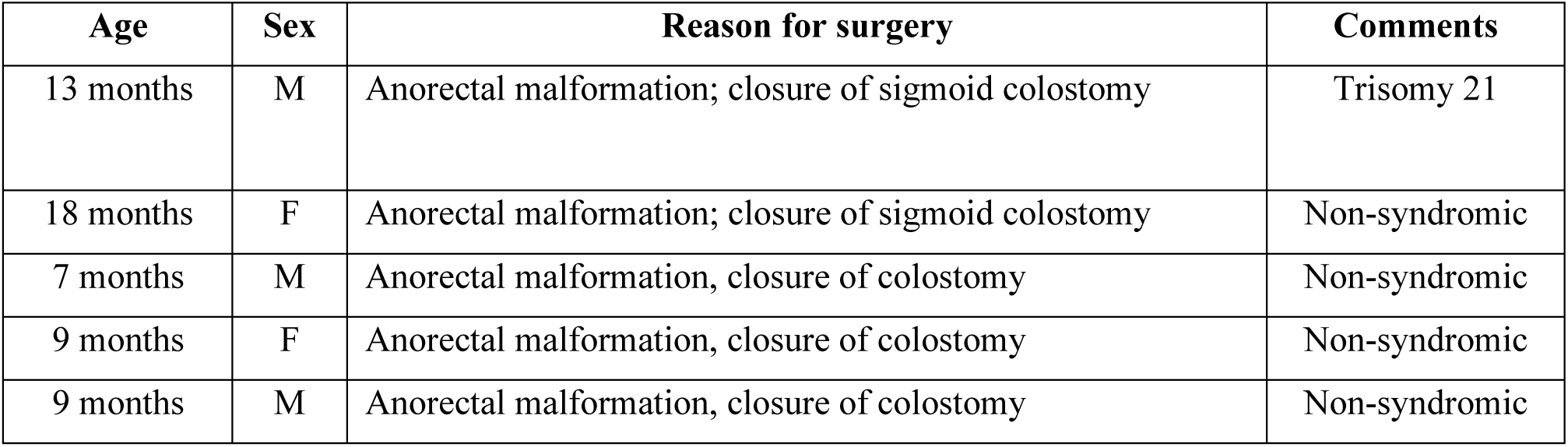
Patient details for tissue preparations used in Ca^2+^ imaging experiments presented in this study.

**Table S2.** Pharmacological tool compounds and indicator dyes used in this study. Preparations were preincubated with antagonists for 20 min at 37°C prior to agonist exposure.

| Compound | Target | Concentration | Supplier |
| --- | --- | --- | --- |
| 43GAP26 | Connexin43 hemichannel blocker | 50 μM | Tocris Bioscience |
| ATP | Purinergic agonist | 100 μM | Sigma-Aldrich |
| Carbenoxolone disodium salt (CBX) | Gap junction blocker | 50 μM | Sigma-Aldrich |
| Cyclopiazonic acid (CPA) | SERCA inhibitor | 30 μM | Abcam |
| Endothelin-1 | ET receptor agonist | 3 μM | Mimotopes |
| Fura2-AM | Ca<sup>2+</sup> indicator dye | 3 μM | Abcam |
| Fluo4-AM | Ca<sup>2+</sup> indicator dye | 4 μM | ThermoFisher |
| Fluo8-AM | Ca<sup>2+</sup> indicator dye | 6 μM | Abcam |
| GR64349 | NK<sub>2</sub>R agonist | 500 nM | Tocris Bioscience |
| GSK1016790A | TRPV4 activator | 100nM; 10 pM to 1μM | Sigma-Aldrich |
| GSK2193874 | TRPV4 blocker | 1 μM | Abcam |
| HC067047 | TRPV4 blocker | 10 μM | Abcam |
| MRS2365 | P2Y<sub>1</sub>R agonist | 250 nM | Tocris Bioscience |
| NKA | NK<sub>2</sub>R agonist | 500 nM | Tocris Bioscience |
| Picrylsulfonic acid (5% solution in H<sub>2</sub>O) | Induces chemical colitis | 2.5% v/v in 50% ethanol | Sigma-Aldrich |
| PPADS | Purinergic antagonist | 50 μM | Abcam |
| Suramin | Purinergic antagonist | 50 μM | Abcam |
| Tetrodotoxin (TTX) | Blocks neurotransmission | 1 μM | Alomone Labs |
| TFLLR-NH<sub>2</sub> | PAR1 agonist | 50 μM | Sigma-Aldrich |

**Table S3.** Details of primary antibodies used in this study.

| Cell type | Target | Host Species | Dilution | Supplier | Clone/Catalog number | RRID/Reference |
| --- | --- | --- | --- | --- | --- | --- |
| Vascular endothelia | CD31 | Rat | 1:400 | BioLegend | MEC13.3 | AB_312909 |
| Enteric Glial Cells | GFAP | Chicken | 1:1000-1:2000 | Abcam | ab4674 | AB_304558 |
|  | S100B | Rabbit | 1:1000 | DAKO | Z031129-2 | AB_2315306 |
|  | Sox10 | Goat | 1:25 | Santa Cruz | sc-17342 | AB_2195374 |
|  | Sox10 | Goat | 1:250 | R&D Systems | AF2864 | AB_442208 |
| Extrinsic terminals | CGRP | Goat | 1:1000 | Abcam | ab36001 | AB_725807 |
|  | DsRed1 | Rabbit | 1:1000 | Clontech | 632496 | AB_10013483 |
|  | Tachykinin | Rat | 1:800 | Abcam | NC1/34 | AB_449518 |
| mMac | CD68 | Rat | 1:1000 | BioLegend | FA.11 | AB_2044004 |
|  | CD169 | Rat | 1:400 | BioLegend | 3D6.112 | AB_10916523 |
|  | F4/80-A488 | Rat | 1:400 | BioLegend | BM8 | AB_893479 |
|  | GFP | Rabbit | 1:1000 | ThermoScientific | A11122 | AB_221569 |
|  | IBA1 | Rabbit | 1:1000 | Wako | 019-19741 | AB_839504 |
|  | MHCII | Rat | 1:500 | BioLegend | M5/114.15.2 | AB_313317 |
| mMac Depletion | CSFR1 (CD115) | Rat | 18.75 µg/g | InVivoMab BioXcell | AFS98 | AB_2687699 |
|  | IgG2A | Rat | 18.75 µg/g | InVivoMab BioXcell | 2A3 | AB_1107769 |
| Lymphatic endothelia | Lyve-1 | Rat | 1:1000 | R&D Systems | MAB2125 | AB_2138528 |
|  | VEGFR3 | Sheep | 1:400 | R&D Systems | AF743 | AB_355563 |
| Neurons | Hu | Human | 1:40000 | - | - | [57] |
|  | HuC/D | Mouse | 1:400 | ThermoScientific | 16A11 | AB_221448 |
|  | NF200 | Mouse | 1:500 | Sigma Aldrich | NE14 | AB_260781 |
|  | PGP9.5 | Rabbit | 1:1000-1:2000 | Abcam | EPR4118 | AB_10891773 |
| PDGFRα<br>cells | PDGFRα | Goat | 1:250 | R&D Systems | AF1062 | AB_2236897 |

**Table S4.** Pair Correlation Function (PCF) data summarizing spatial connectivity of enteric glial Ca^2+^ signaling to GSK101 (100nM) under different conditions and at different distances.

**PCF at 60 s**
| **T=60 s** |  |  |  |  |  |  |  |  |  |  |
| --- | --- | --- | --- | --- | --- | --- | --- | --- | --- | --- |
| **Distance**<br>**(microns)** | **Control** |  | **CBX** |  | **SUR+PPADS** |  | **CPA** |  | **GAP26** |  |
|  | **Mean** | **(95%**<br>**CI)** | **Mean** | **(95%**<br>**CI)** | **Mean** | **(95%**<br>**CI)** | **Mean** | **(95%**<br>**CI)** | **Mean** | **(95%**<br>**CI)** |
| **58** | 0.944 | 0.216 | 0.975 | 0.062 | 1.040 | 0.014 | 1.006 | 0.032 | 1.010 | 0.115 |
| **115** | 1.278 | 0.550 | 0.966 | 0.071 | 1.021 | 0.046 | 0.997 | 0.024 | 1.075 | 0.053 |
| **173** | 0.874 | 0.192 | 0.976 | 0.148 | 0.980 | 0.060 | 0.953 | 0.032 | 1.007 | 0.165 |
| **230** | 1.101 | 0.289 | 0.997 | 0.067 | 0.949 | 0.073 | 0.976 | 0.023 | 1.005 | 0.054 |
| **289** | 0.893 | 0.199 | 1.015 | 0.084 | 1.104 | 0.105 | 0.987 | 0.027 | 0.948 | 0.181 |
| **346** | 0.922 | 0.210 | 1.079 | 0.138 | 0.984 | 0.068 | 0.973 | 0.029 | 1.010 | 0.072 |

**TNBS**
| **T=30 s** |  |  |  |  | **T=60 s** |  |  |  |
| --- | --- | --- | --- | --- | --- | --- | --- | --- |
| **Distance**<br>**(microns)** | **Control** |  | **TNBS** |  | **Control** |  | **TNBS** |  |
|  | **Mean** | **(95%**<br>**CI)** | **Mean** | **(95%**<br>**CI)** | **Mean** | **(95%**<br>**CI)** | **Mean** | **(95%**<br>**CI)** |
| **58** | 0.974 | 0.275 | 1.020 | 0.038 | 0.860 | 0.162 | 1.028 | 0.011 |
| **115** | 1.444 | 0.766 | 1.060 | 0.064 | 1.268 | 0.460 | 1.036 | 0.034 |
| **173** | 0.855 | 0.285 | 1.030 | 0.074 | 0.923 | 0.171 | 1.027 | 0.043 |
| **230** | 0.857 | 0.213 | 0.999 | 0.023 | 1.078 | 0.241 | 1.008 | 0.025 |
| **289** | 0.930 | 0.202 | 0.924 | 0.049 | 0.844 | 0.171 | 0.959 | 0.049 |
| **346** | 0.982 | 0.366 | 1.020 | 0.029 | 0.885 | 0.177 | 0.991 | 0.017 |

**TTX**
| **T=30 s** |  |  |  |  | **T=60 s** |  |  |  |
| --- | --- | --- | --- | --- | --- | --- | --- | --- |
| **Distance**<br>**(microns)** | **Control** |  | **TTX** |  | **Control** |  | **TTX** |  |
|  | **Mean** | **(95%**<br>**CI)** | **Mean** | **(95%**<br>**CI)** | **Mean** | **(95%**<br>**CI)** | **Mean** | **(95%**<br>**CI)** |
| **58** | 1.390 | 0.716 | 1.032 | 0.120 | 1.094 | 0.192 | 1.019 | 0.090 |
| **115** | 0.937 | 0.072 | 0.961 | 0.173 | 0.944 | 0.116 | 0.963 | 0.072 |
| **173** | 0.984 | 0.292 | 0.799 | 0.082 | 1.053 | 0.158 | 0.972 | 0.122 |
| **230** | 1.017 | 0.491 | 1.137 | 0.342 | 1.030 | 0.197 | 1.119 | 0.331 |
| **289** | 0.761 | 0.358 | 0.758 | 0.306 | 0.888 | 0.230 | 1.015 | 0.116 |
| **346** | 1.066 | 0.811 | 1.004 | 0.148 | 1.042 | 0.258 | 0.959 | 0.111 |

## SUPPLEMENTARY FIGURES

**Figure S1.**
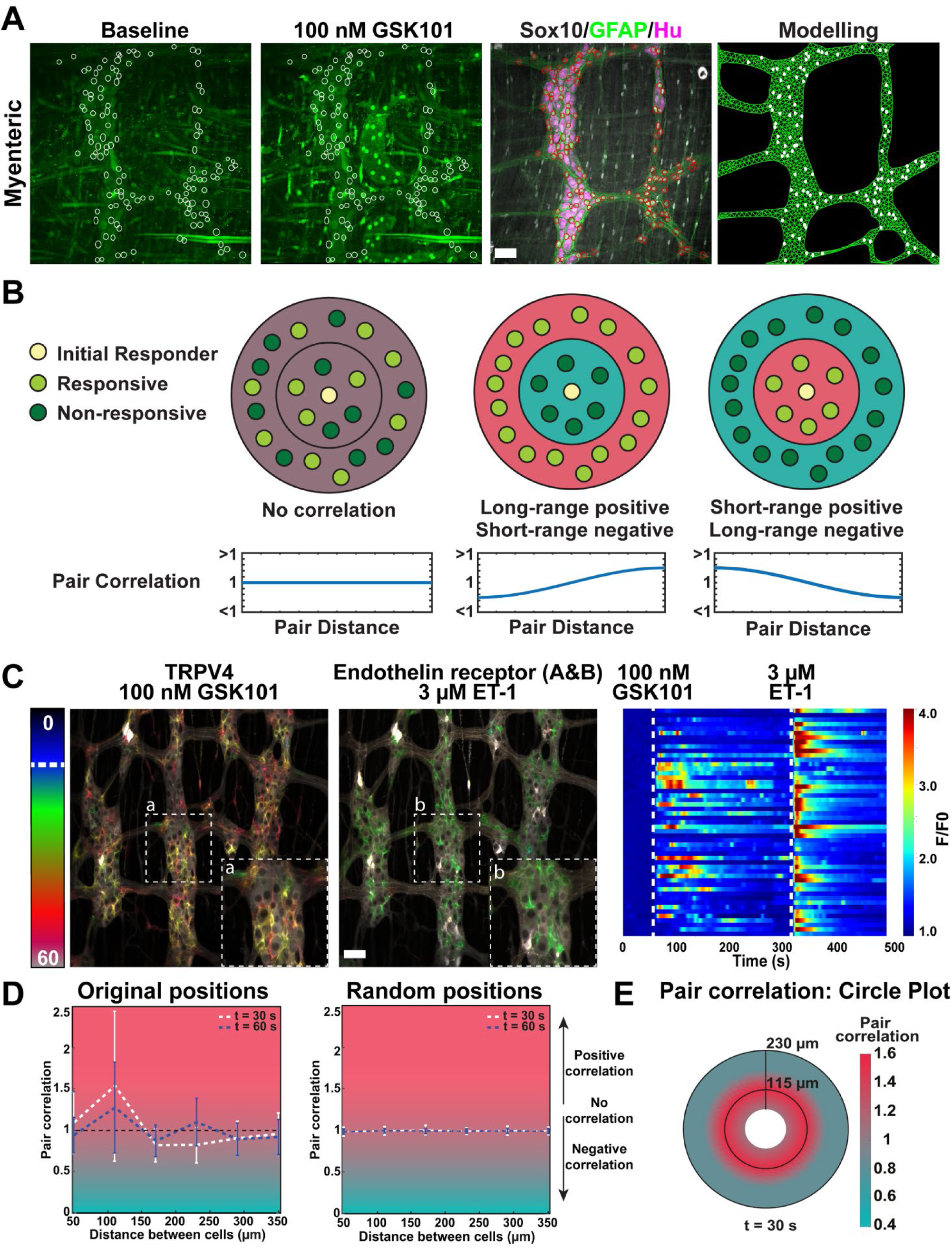
Enteric glial cells display varying Ca^2+^ kinetics upon activation of different receptors, and function as a coordinated network in the myenteric plexus. **A.** The coordinates for each glial cell ROI (white circles), the corresponding GSK101-mediated Ca^2+^ responses, and the coordinates of the ganglia were extracted for each experiment. The coordinates corresponding to the outlines of ganglia were used to calculate the distance and adjacency matrix (‘Modelling’; Scale bar: 100 µm). **B.** The extracted spatiotemporal information was used to calculate PCFs. These are represented as circle plots, where the radius of the circle is proportional to the pair distances between cells. If a cell responds first to a stimulus (initial responder: light yellow), and activates surrounding cells in a random fashion, then the PCF will be 1 at all pair distances. However, if the initial responder activates cells further away (responsive: light green), then this leads to long-range positive (red shading at larger distances) and short-range negative PCF (teal shading for smaller distances). If apposing cells are activated by an initial responder, then the correlation is short-range positive (red shading) and long-range negative (teal shading). **C.** The application of GSK101 (100 nM) resulted in varying temporal Ca^2+^ dynamics across the myenteric glial network. The Ca^2+^ waves propagated from early responders (color scale= green and green-yellow) to surrounding glial cells (color scale: yellow and red; **inset a**). However, the subsequent activation of endothelin receptors (Endothelin-1, 3 µM) resulted in an instantaneous and uniform Ca^2+^ response by glia throughout the ganglia (green; **inset b**). Each image is a 60 s temporal color-coded projection, where the white dotted line on the LUT represents the time of drug application (10 s; Scale bar = 50 µm). The kinetics of the GSK101 and ET-1 evoked Ca^2+^ responses are also shown as a heatmap. **D.** The pair correlation functions (PCFs) were calculated to examine the coordinated activity of glia in response to GSK101. PCFs were calculated between pairs of glial cells at varying distances at t=30 s and 60 s after GSK101 addition. The PCF was the highest for glial cells separated by 115 µm at t=30 s, indicating highly coordinated activity. When the positions of the glial cells were shuffled randomly 100 times, the average PCF value was calculated to be 1 at all pair distances, indicating that the coordinated activity was abolished. **E.** PCF can be better visualized as a circle plot, where the radius of the plot is the pair distance between glia, and the color-coding indicates the degree of correlated activity (red is high and green is low PCF). As the highest correlation was at t=30 s, this timepoint was chosen for further analysis (n=9; 1059 cells).

**Figure S2.**
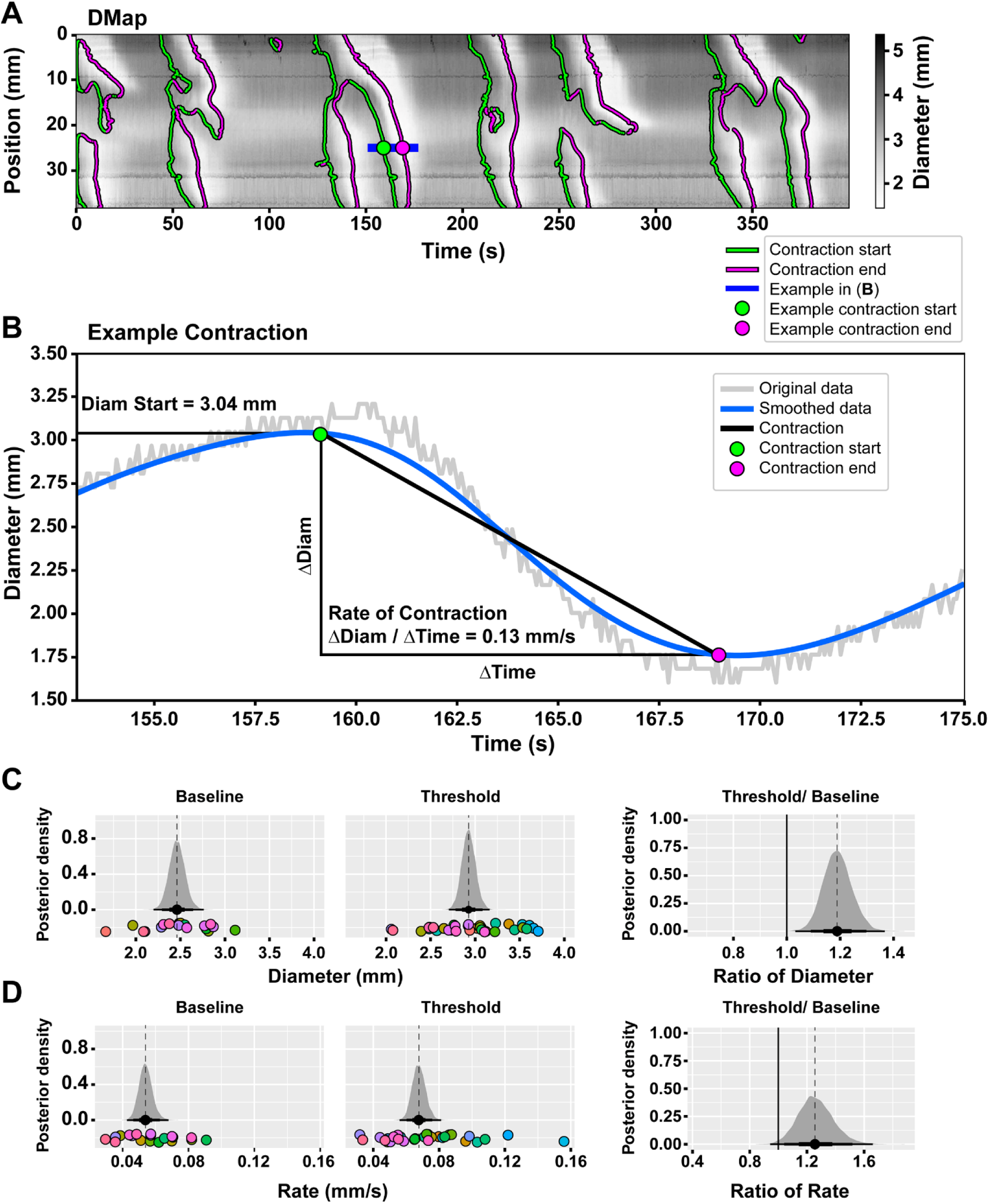
Analytical method used to study diameter maps of colonic motility. **A.** A representative spatiotemporal map showing changes in diameter along the length of a mouse colon over time. Overlaid colored lines demonstrate how the software identifies the start (green) and end (magenta) of contractions. The blue horizontal line located between 150-200s indicates an example contraction. The contraction starts at a specified location and time on the recording (green circle), The event progresses over time at the same location until the diameter no longer narrows. This spatiotemporal location defines the contraction end (magenta circle). All contractions captured in the map are included in this analysis, including those not classified as CMCs. An event, such as a CMC, is considered to consist of multiple contractions across multiple spatiotemporal locations. For example, this image contains a total of 3907 discrete contractions over 478 spatial locations (resolution of 478/38 mm = 12.5 spatial locations per mm), with an average of 3907/478 = 8.2 contractions per spatial position. **B.** The highlighted contraction in ***A*** (blue line) is expanded as an XY subplot (note that the Y axis is gut diameter, not position). The rate of contraction is given by the absolute slope (Δ diameter / Δ time; black line), where a contraction is detected by the interval where the smoothed data gradient is below the threshold of -0.02 mm/s (i.e., a negative gradient). The example shows a rate of contraction of 0.13 mm/s, with a diameter start of 3.04 mm. All contractions were averaged to produce a single number for both the *mean rate of contractions* (MRC) and the *mean diameter of contractions* (MDC) in each recording. Posterior distribution graphs demonstrate the probability of a contraction occurring at any given diameter (**C**) or occurring at any given rate (**D**). As an example, posterior distribution graphs in ***C*** and ***D*** show these probabilities when tissues were at baseline (*left*) or at threshold (*right*) intraluminal pressures. Individual color dots represent mean data points from each individual experiment. The posterior distribution of the mean values is represented by the grey shading and dashed vertical lines indicate the mean. The ruler bar at the base of each posterior distribution highlights the middle 68, 95 and 99.7 percentiles. To assess differences in these distributions, the mean values between conditions were compared as a ratio (*right panels*). The solid vertical line on these ratio graphs indicates where the ratio is 1. In cases where the vertical line is positioned outside the 95% posterior this depicts a real effect (thinnest line on ruler bar). Where the vertical line is positioned inside the 95% posterior, this indicates that there was no detectable difference (two thicker lines on the ruler bar). **C.** The MDC at baseline ranged from 2.29-2.64 mm vs. 2.60-2.97 mm at threshold. The ratio of MDC was 1.13 indicating that colon diameters were greater at threshold. **D.** No difference in the MRC was detected when preparations were stimulated at threshold versus baseline conditions (ratio of the mean=1.10). This example demonstrates a robust, detectable change following a well-established stimulus of colonic contractions.

**Figure S3.**
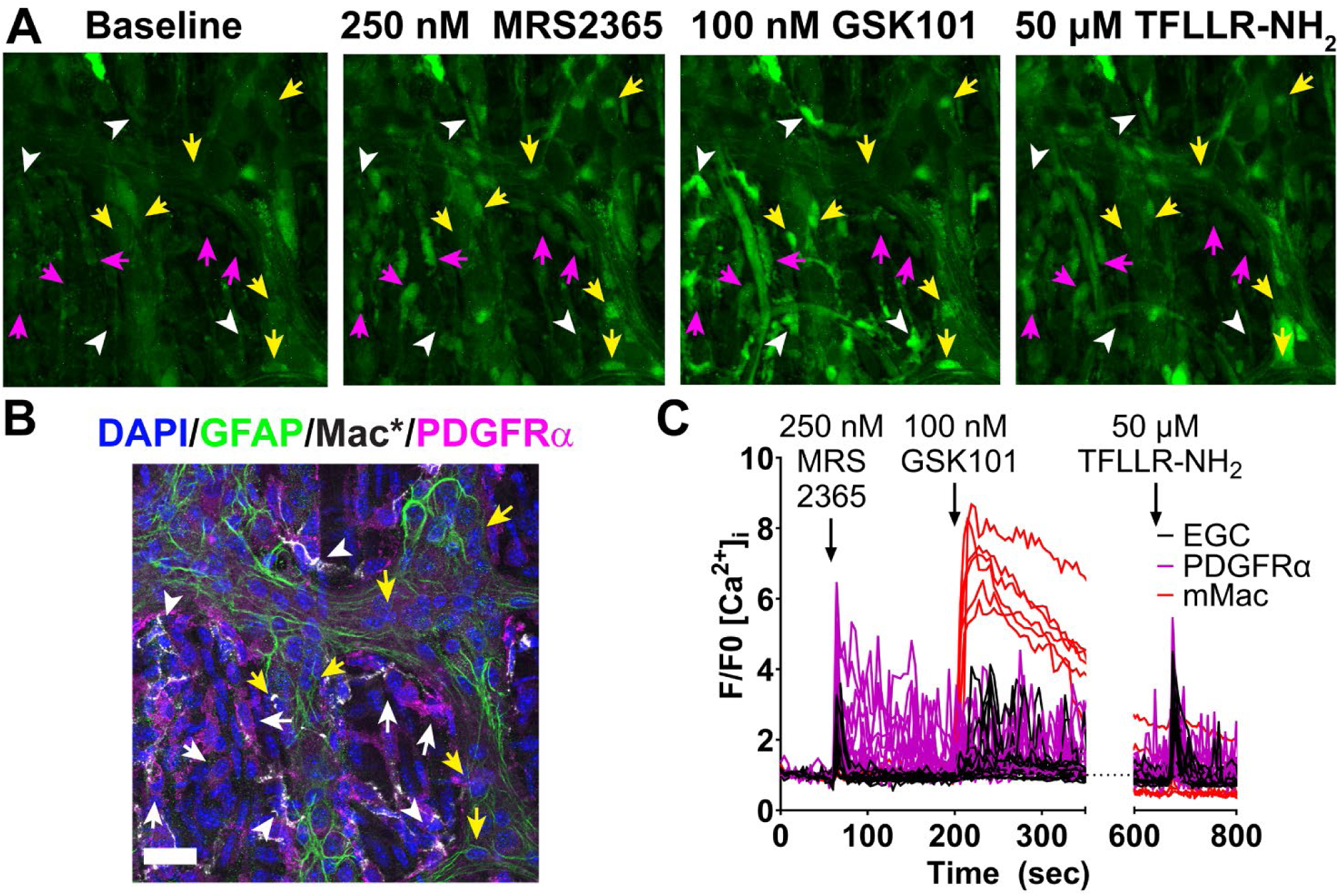
PDGFRα+ cells do not express TRPV4. **A.** PDGFRα+ cells were identified pharmacologically based on their responsiveness to P2Y_1_R (MRS2365) and PAR1 (TFLLR-NH_2_) agonists combined with immunolabeling (**B**, white arrows, PDGFRα-positive: magenta). **C.** Traces from individual cells demonstrate robust Ca^2+^ signals to GSK101 by mMac (white arrowheads) and glia (yellow arrows; GFAP-positive: green), whereas PDGFRα cells in the same preparation did not respond. Scale: 30 µm.

**Figure S4.**
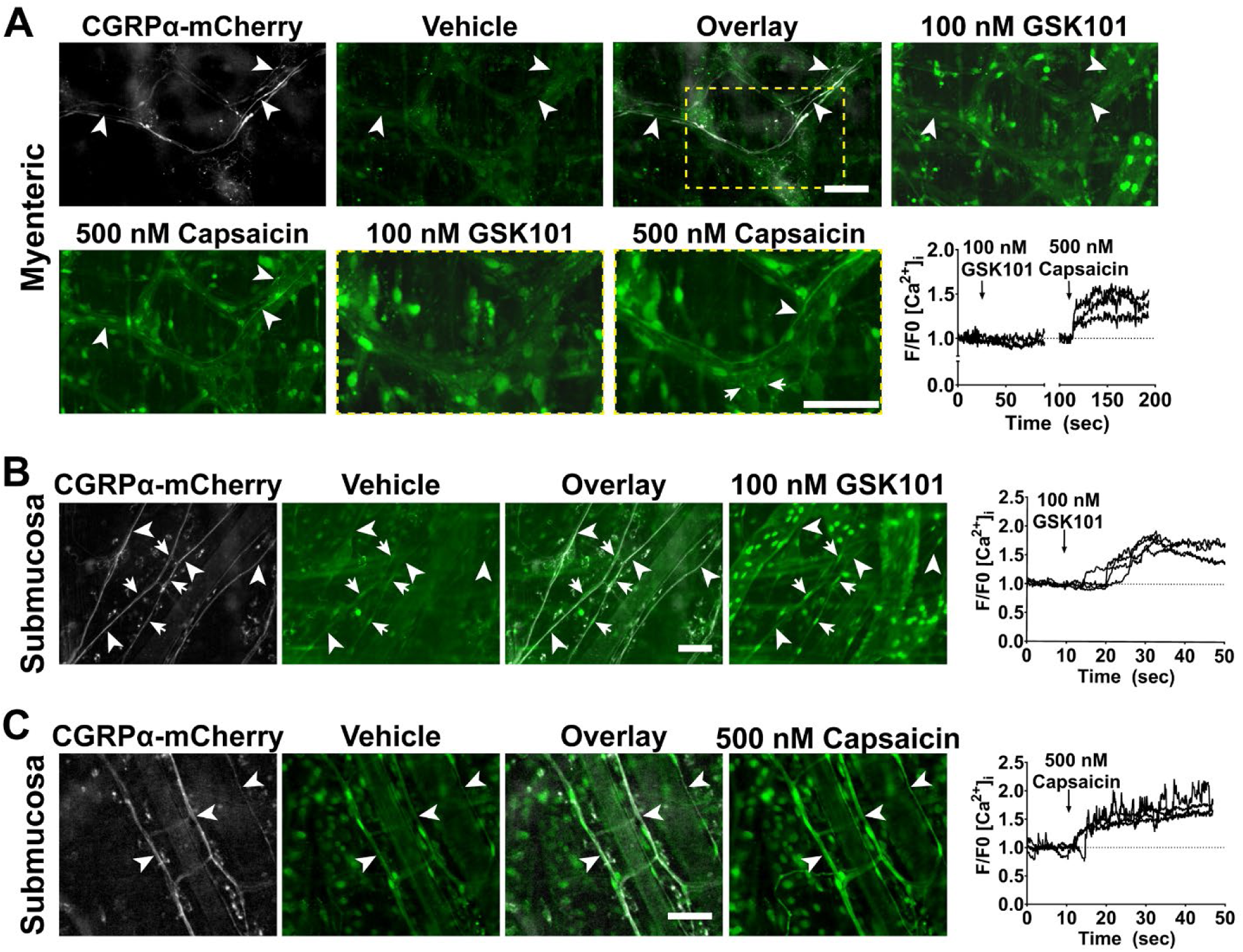
TRPV4 is functionally expressed by a subset of extrinsic sensory afferents innervating the mouse colon. **A.** Extrinsic afferents innervating the CGRPα-mCherry mouse colon were identified by mCherry fluorescence (arrowheads). Afferents within the myenteric region responded to the TRPV1 activator capsaicin, but not to GSK101. Zoomed in views of the area within the yellow rectangle are presented as insets. Responses to capsaicin by extrinsic nerves (arrowhead) and varicosities (arrows) are indicated. **B.** GSK101 evoked Ca^2+^ transients in extrinsic afferent fibers of the submucosal layer (arrowheads) which were mainly associated with activation of associated glial-like cells (arrows). **C.** Capsaicin also activated extrinsic afferents in the submucosa (arrowheads). Scale: 20 µm. Preparations from N=5 mice.

**Figure S5.**
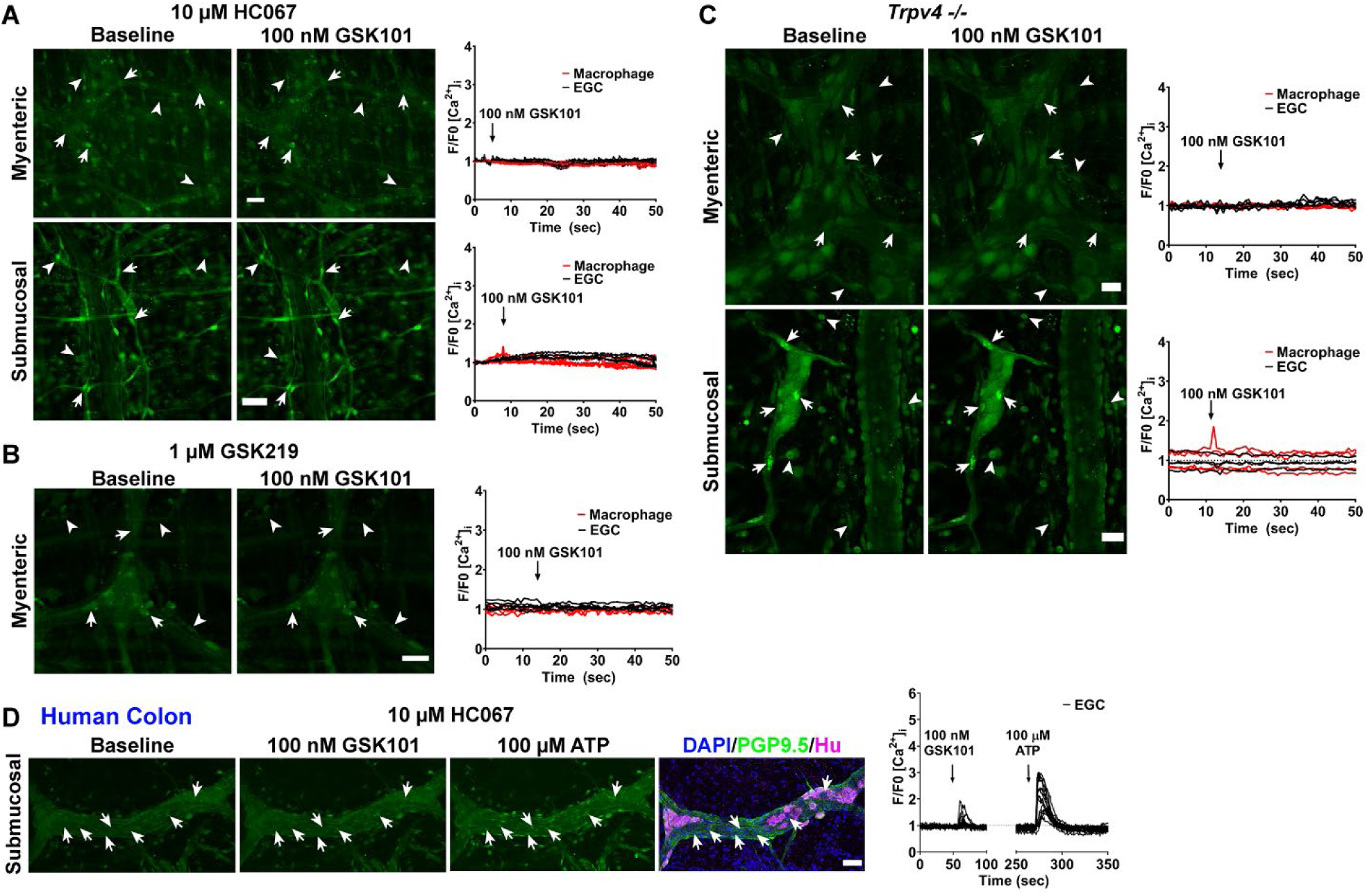
Ca^2+^ responses to GSK1016790A are highly TRPV4-specific. **A-B.** Pre-treatment with the TRPV4 antagonists HC067 (10 µM, n=7, **A**) or GSK219 (1 µM, n=3, **B**) prevented GSK101-evoked Ca^2+^ transients in both the myenteric and submucosal layers of the mouse colon. **C.** No Ca^2+^ responses to GSK101 were observed in tissues from *trpv4*^-/-^ mice (n=9). **D.** GSK101-evoked elevations in Ca^2+^ in the submucosal layer of the human colon were prevented by HC067 (n=4). ATP-evoked responses were retained in these preparations. Scale: 20 µm.

**Figure S6.**
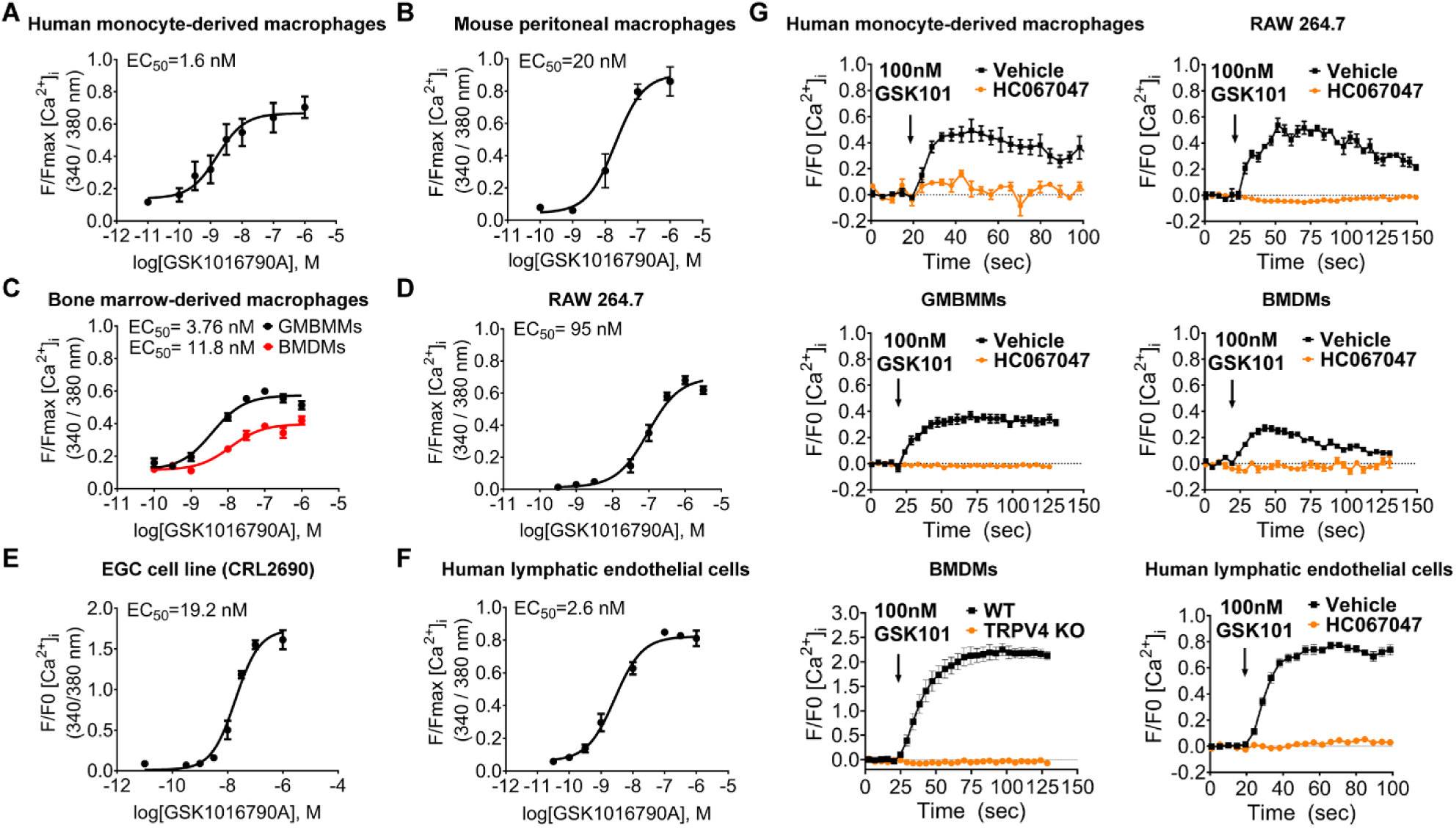
TRPV4 is expressed by macrophages, enteric glia, and lymphatic endothelial cells. Concentration-dependent Ca^2+^ responses to GSK101 were detected in primary human monocyte-derived (**A**), mouse peritoneal (**B**) and GM-CSF- and M-CSF-generated bone marrow-derived macrophages (BMDM, **C**), and in the murine macrophage cell line RAW264.7 (**D**). Functional expression of TRPV4 by enteric glia and lymphatic endothelial cells was demonstrated using the enteric glial cell line CRL2690 (**E**) and primary human dermal lymphatic endothelial cells (**F**). **G.** Effects of GSK101 (100 nM) in all cell types were abolished by the TRPV4-selective antagonist HC067 (10 µM) and were absent in cells derived from TRPV4^-/-^ mice.

**Figure S7.**
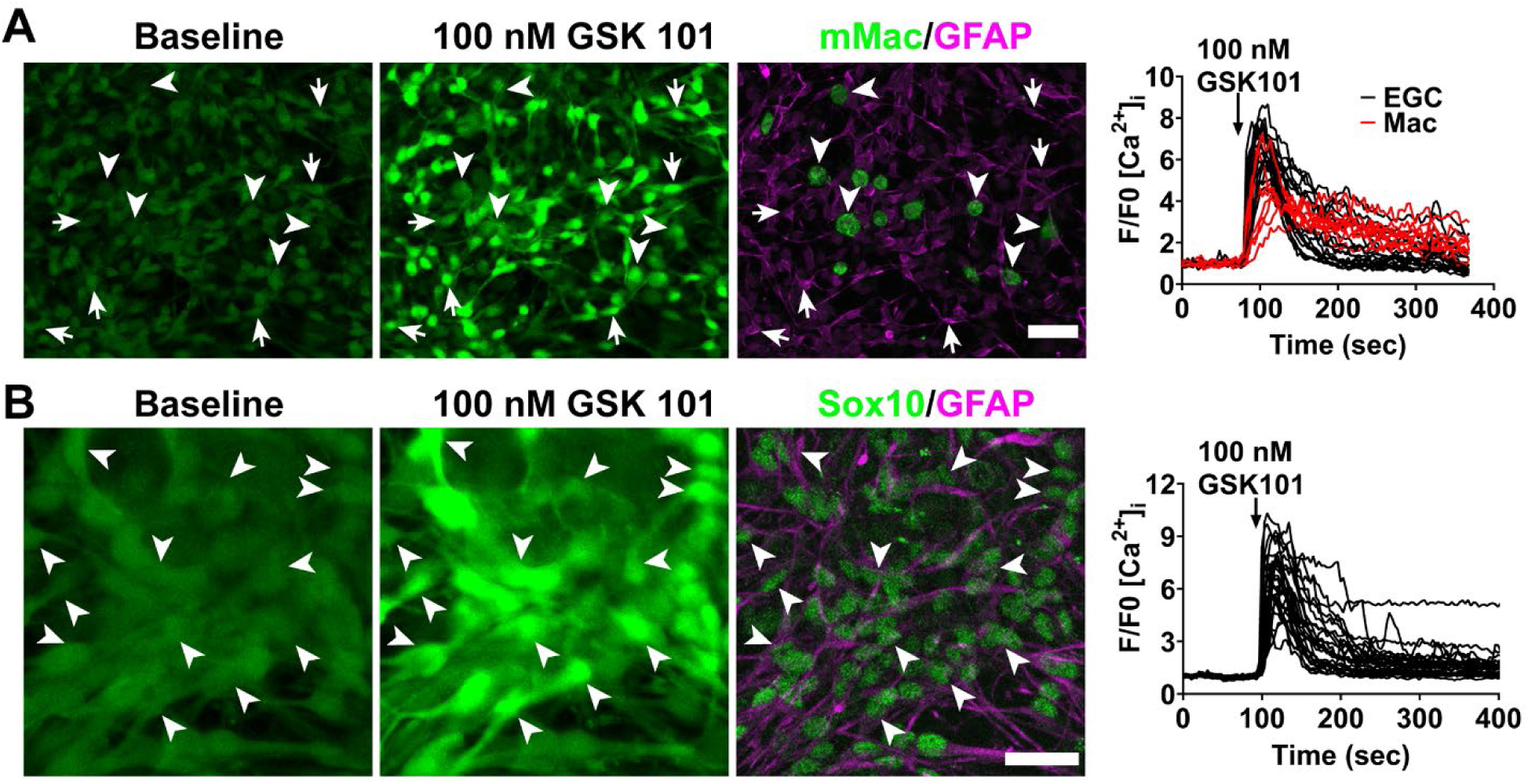
Cultured enteric glia and muscularis macrophages respond directly to TRPV4 activation. **A.** Sustained Ca^2+^ responses to GSK101 (100 nM) were detected in mMac (arrowheads, green) and glia (arrows, magenta) under culture conditions. **B.** Enriched glial cultures exhibited similar responses to GSK101. TRPV4-evoked Ca^2+^ oscillations by cultured glia were not observed, in marked contrast to glia in tissue preparations. Scale: 50 µm.

**Figure S8.**
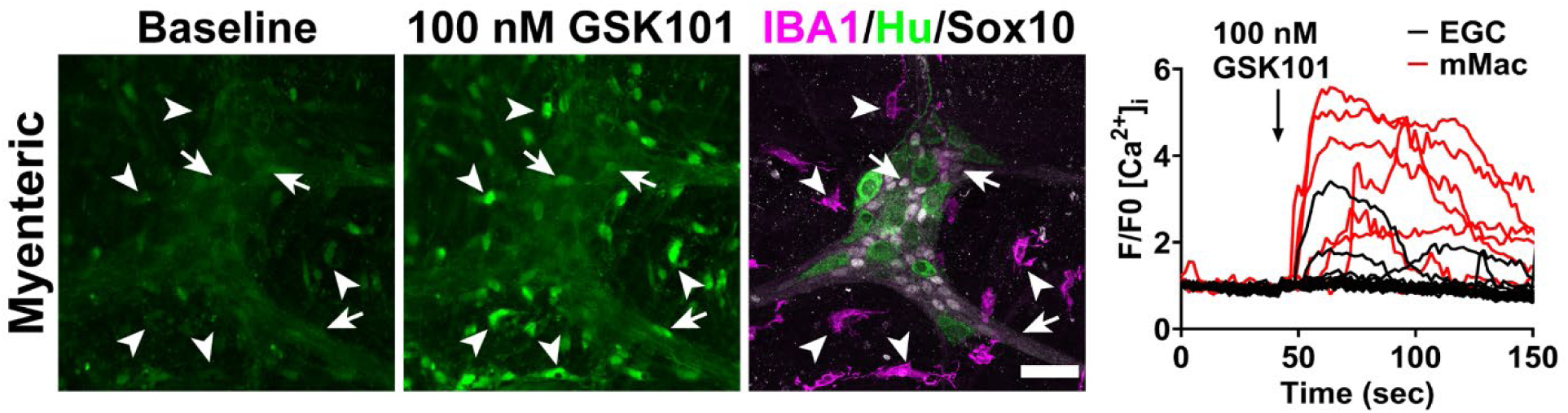
TRPV4 is functionally expressed by enteric glia and macrophages in the monkey colon. Enteric glia (arrows) and mMac (arrowheads) in the myenteric region of the monkey colon exhibited calcium responses to GSK101 (100nM). The identity of glia (Sox10+: white), mMac (IBA1+: magenta), and neurons (Hu+: green) were confirmed by immunostaining. Scale = 50 µm.

**Figure S9.**
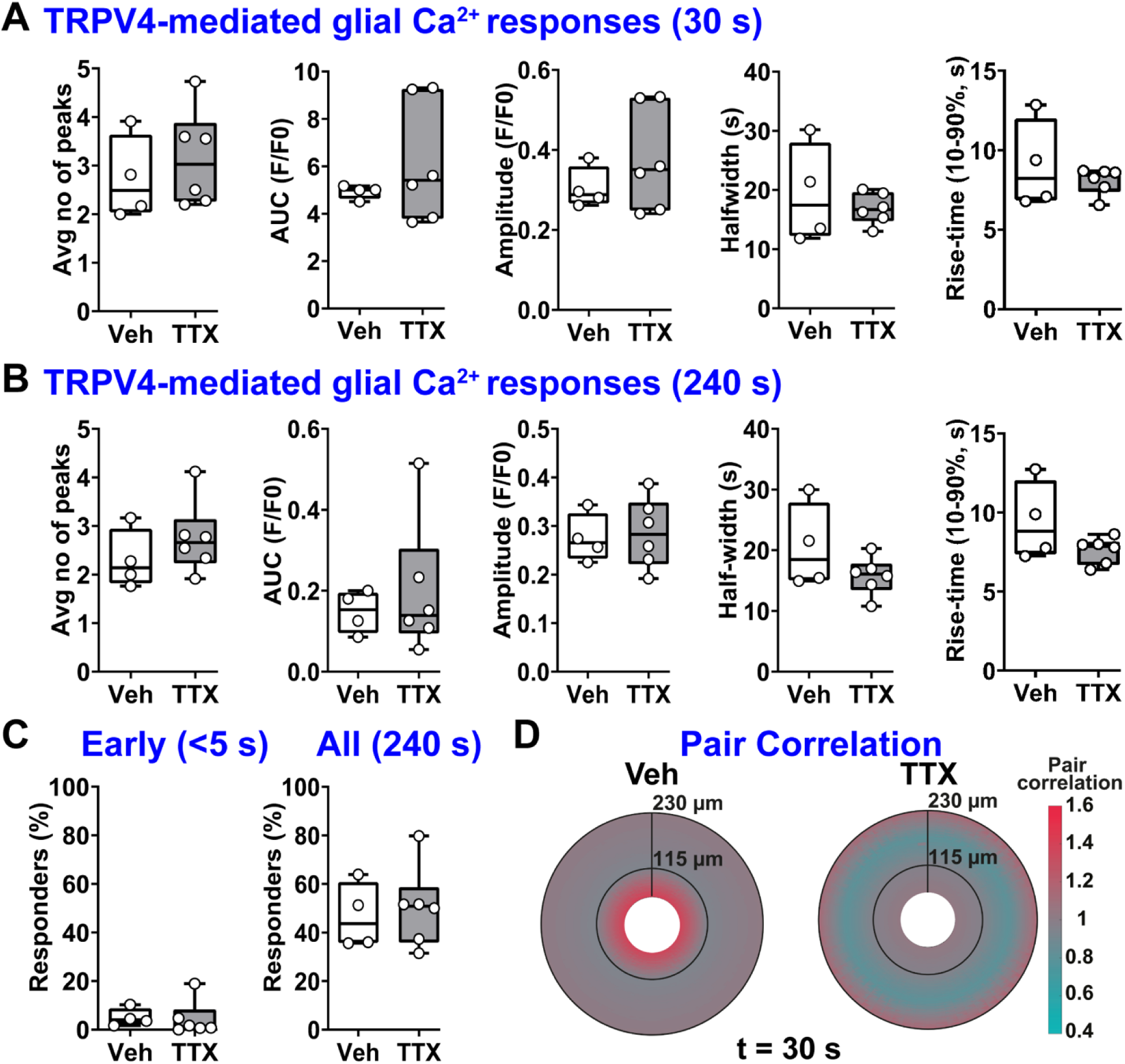
TRPV4-dependent activation of the glial network is largely independent of neuronal influence. Block of neurotransmission (TTX, 100 nM) did not significantly alter the magnitude and nature of glial responses to GSK101 (100 nM) at early (30s, **A**) or later (240s, **B**) time points. **C.** TTX did not affect the relative number or proportion of early and late responding glia. **D.** PCF analysis performed using data at 30s identified reduced coordination of glial responses in the presence of TTX at pair distances of 50 µm. Data are expressed as mean ± 95% C.I. The horizontal line in the graph represents the mean, and whiskers denote the minimum and maximum values. Groups were compared by one-way ANOVA with Dunnett’s post hoc test.

**Figure S10.**
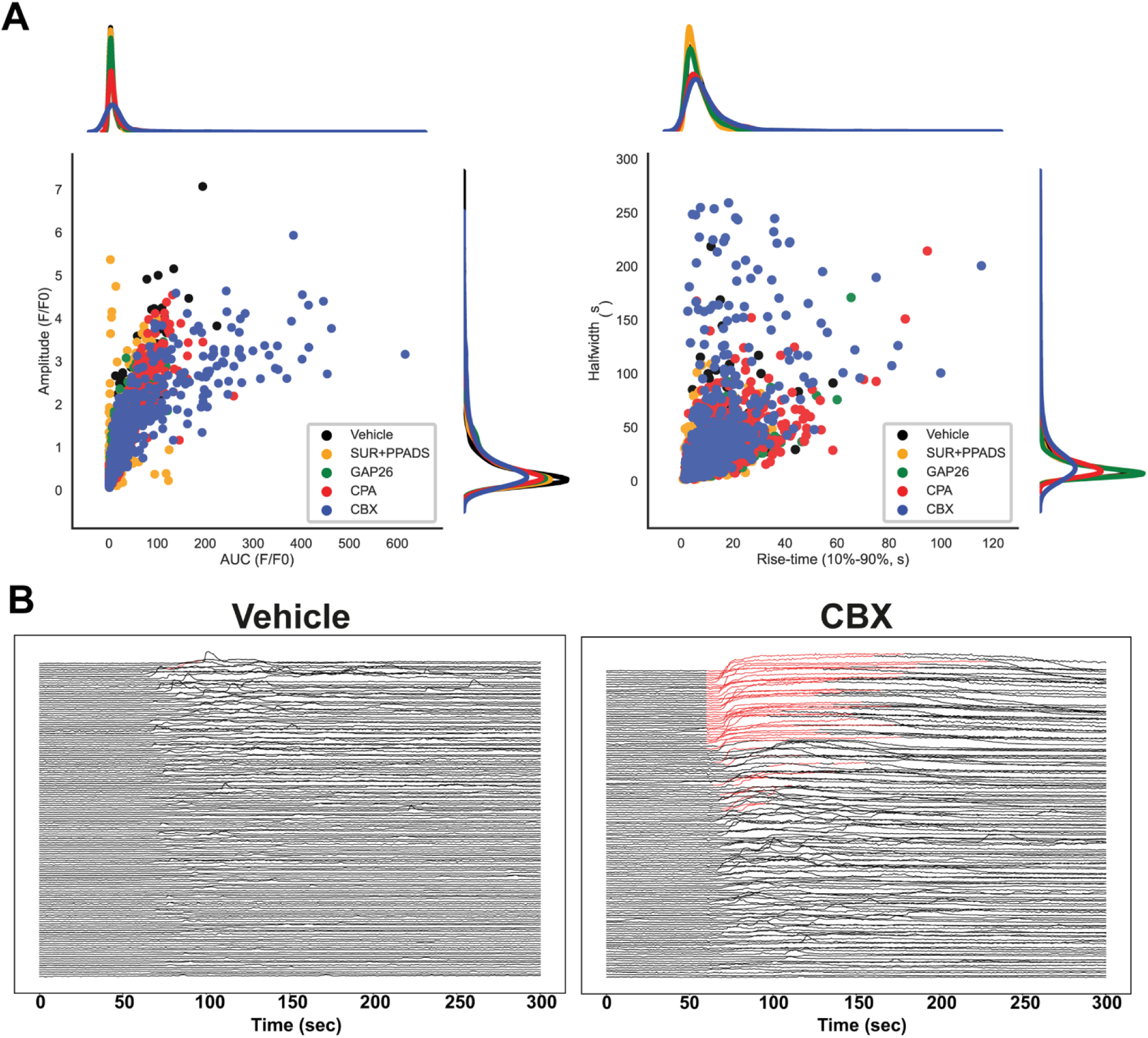
Summary of mechanistic studies to determine how TRPV4 promotes sustained glial activity. **A.** Scatter plot with kernel density estimation (KDE) to visualize the distribution of TRPV4-mediated glial Ca^2+^ responses. Pretreatment with CBX resulted in increased amplitude, AUC, and rise-time for Ca^2+^ transients in glia. **B**. Representative example comparing GSK101-evoked Ca^2+^ signaling in glia following treatment with vehicle or CBX. Glia exhibiting a sustained response (AUC>150) are colored in red.

**Figure S11.**
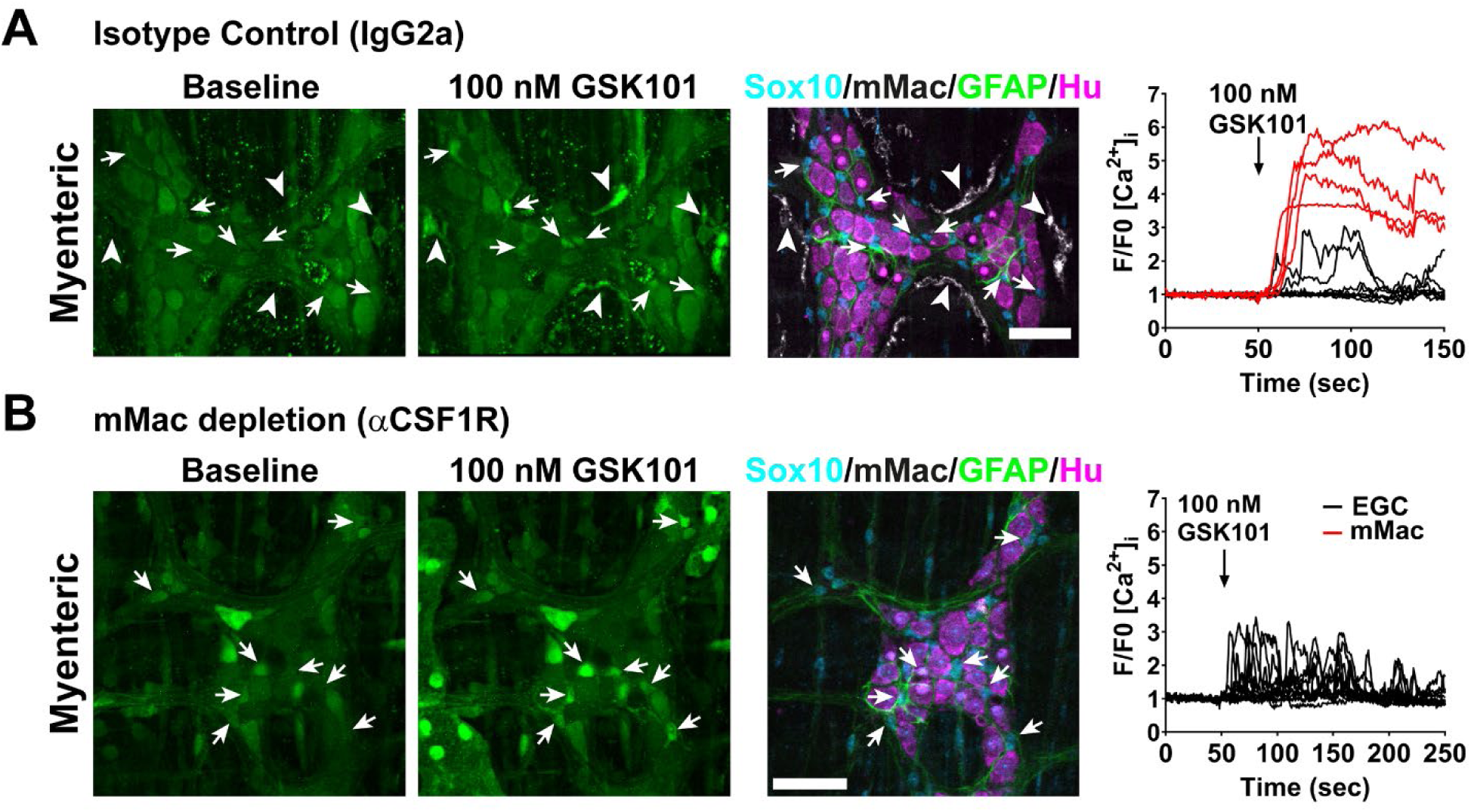
TRPV4-dependent activation of enteric glial cells does not require mMac. **A.** Responses by myenteric glia (arrows) and mMac (arrowheads) to GSK101 (100 nM) were retained in colonic wholemounts from IgG2a-treated mice (isotype control, n=4). **B.** Acute mMac depletion (αCSF1R mAb, n=4) did not impact Ca^2+^ responses by glia (white arrows) to GSK101. These data are consistent with the direct activation of these cells and provide further support for expression of TRPV4 by glia. Scale: 50 µm.

**Figure S12.**
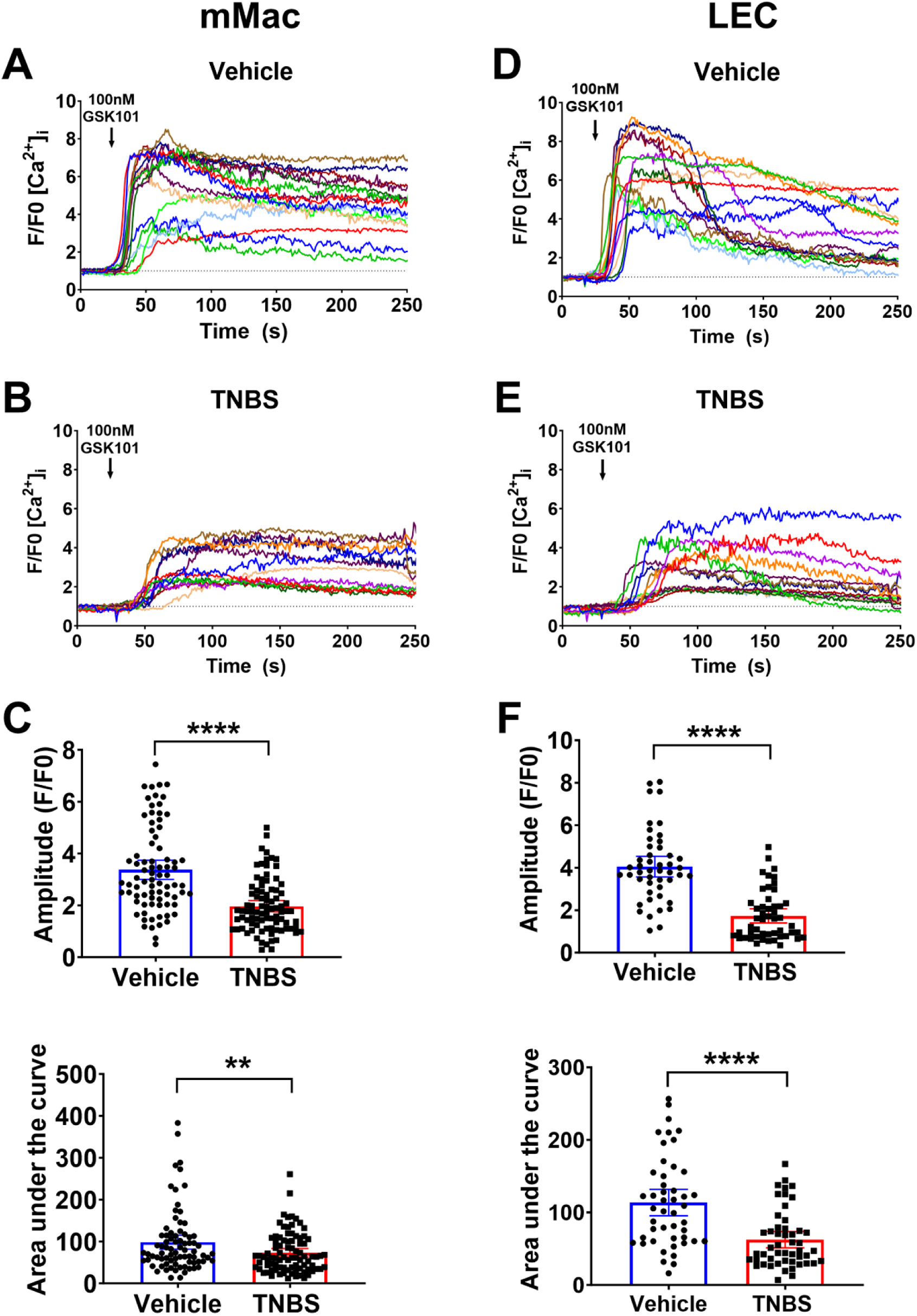
Inflammation-associated suppression of TRPV4-dependent Ca^2+^ signaling in mMac and LECs. Acute TNBS colitis was associated with a suppression of TRPV4-dependent Ca^2+^ signaling in mMac (**A, B;** healthy: n=79 mMac, N=5, TNBS: n=87 mMac, N=6) and LEC (**D, E;** healthy: n=47 LEC, N=5, TNBS: n=50 LEC, N=6). This reduction was reflected by a reduced amplitude of Ca^2+^ peaks and by a significantly smaller area under the curve (**C, F**). Each trace and data point represents Ca^2+^ signaling recorded from an individual cell. Data are expressed as mean ± 95% C.I., **, p<0.01, ****, p<0.0001, unpaired t-test.

**Figure S13.**
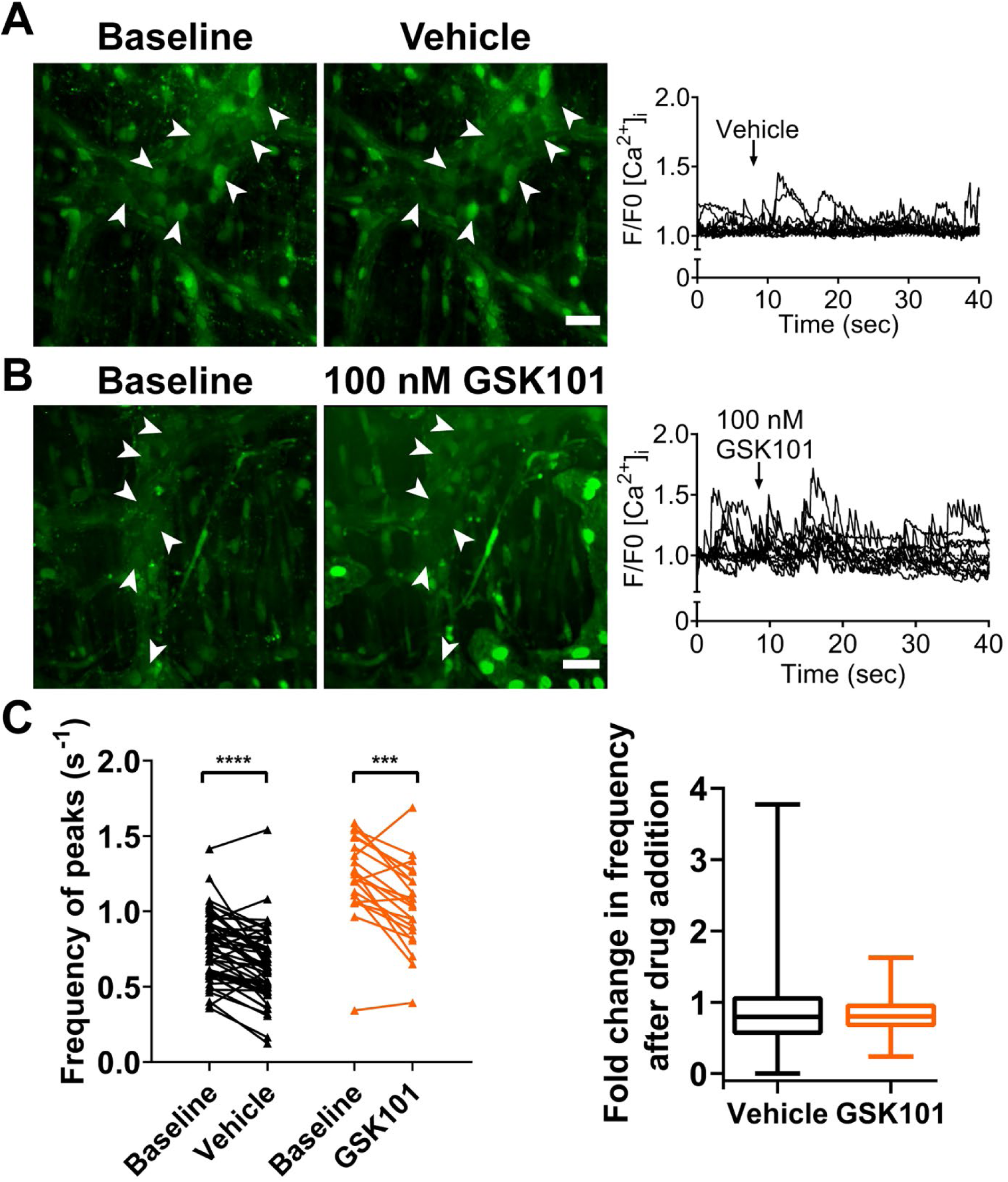
TRPV4 activation does not affect spontaneous activity of myenteric neurons. Myenteric wholemounts of the mouse colon were imaged at 70 fps to determine the effect of TRPV4 activation (GSK101, 100nM) on spontaneous Ca^2+^ transients. Neither vehicle (**A**) nor GSK101 (**B**) treatment promoted increased Ca^2+^ signaling. **C.** There was a significant reduction in activity over time in both vehicle and GSK101 treated groups. The horizontal line in each graph represents the mean, and whiskers denote the minimum and maximum values. N=3 for GSK101 and n=4 for vehicle.

**Figure S14.**
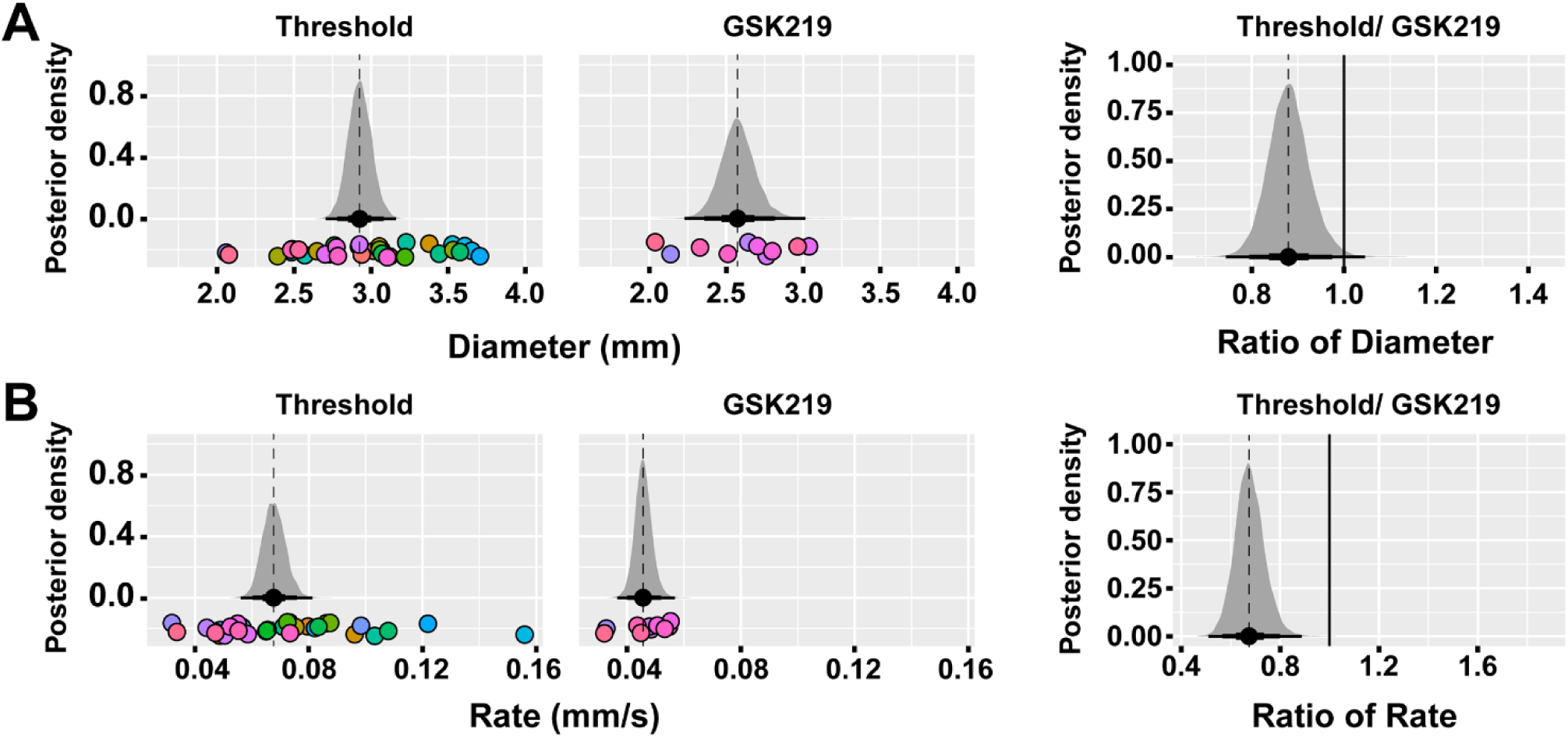
Pharmacological inhibition of TRPV4 reduces MRC in the isolated mouse colon. **A**. Probability distribution graphs demonstrate the likelihood of contractions occurring at different mean colon diameters in control (‘*threshold*’, left) or GSK219-treated (middle) conditions. No difference in the mean diameter of contractions was detected between groups (ratio of mean=0.93). **B**. The mean rate of contraction was reduced in the presence of GSK219 compared to control conditions (ratio of mean=0.8). Each colored dot represents an individual experiment (control: n=34, GSK219: n=10 mice).

**Figure S15.**
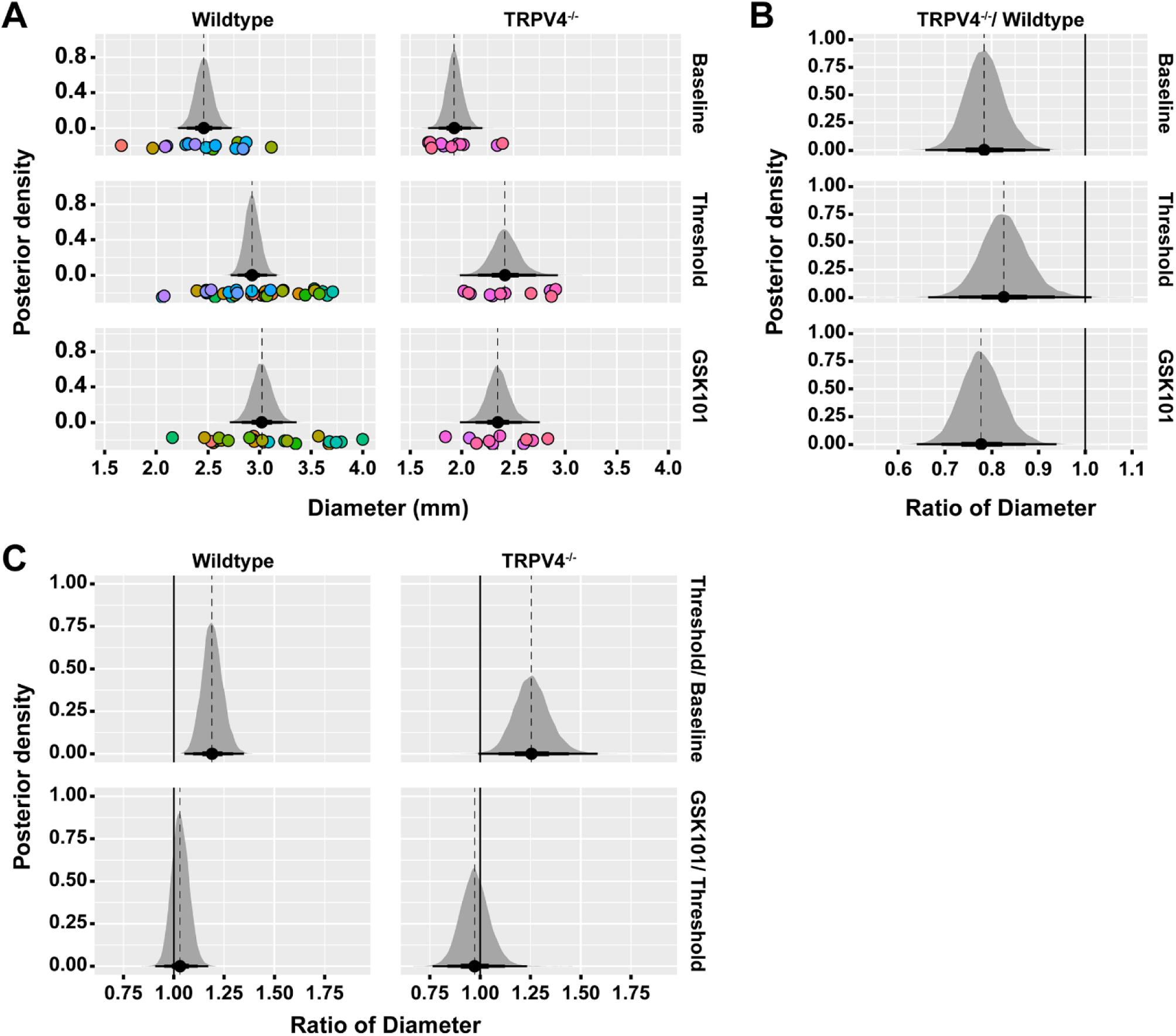
Deletion of TRPV4 reduces the colon diameter of a contraction. **A.** Probability distribution graphs demonstrate the mean diameter of contractions (MDC) for wildtype (*left*) and TRPV4^-/-^ (*right*) colons under baseline, threshold, and GSK101-treated conditions (*top-bottom*). **B.** The ratio of MDCs was compared between wildtype and TRPV4^-/-^ colons at baseline (n=20 and n=11, *top*), threshold (n=34, n=11, *middle*) and GSK101-treated conditions (n=24 and n=11, *bottom*). The MDC of contractions was lower in TRPV4^-/-^ tissue compared to wildtype under all conditions tested (ratio of mean-baseline= 0.78, threshold= 0.87, GSK101= 0.82). **C.** Separate comparisons were made for the ratio of MDC at threshold/baseline (*top*) and threshold/GSK101 (*bottom*) for wildtype (*left*) and TRPV4^-/-^ colons (*right*), respectively. The MDC was greater at threshold compared to baseline for wildtype and TRPV4^-/-^ tissues (ratio of mean: wildtype= 1.13, TRPV4^-/-^ = 1.25). There was no difference in the MDC at threshold when compared to GSK101-treated (ratio of mean: WT= 1.02, TRPV4^-/-^= 0.97). Each colored dot represents an individual experiment.

**Figure S16.**
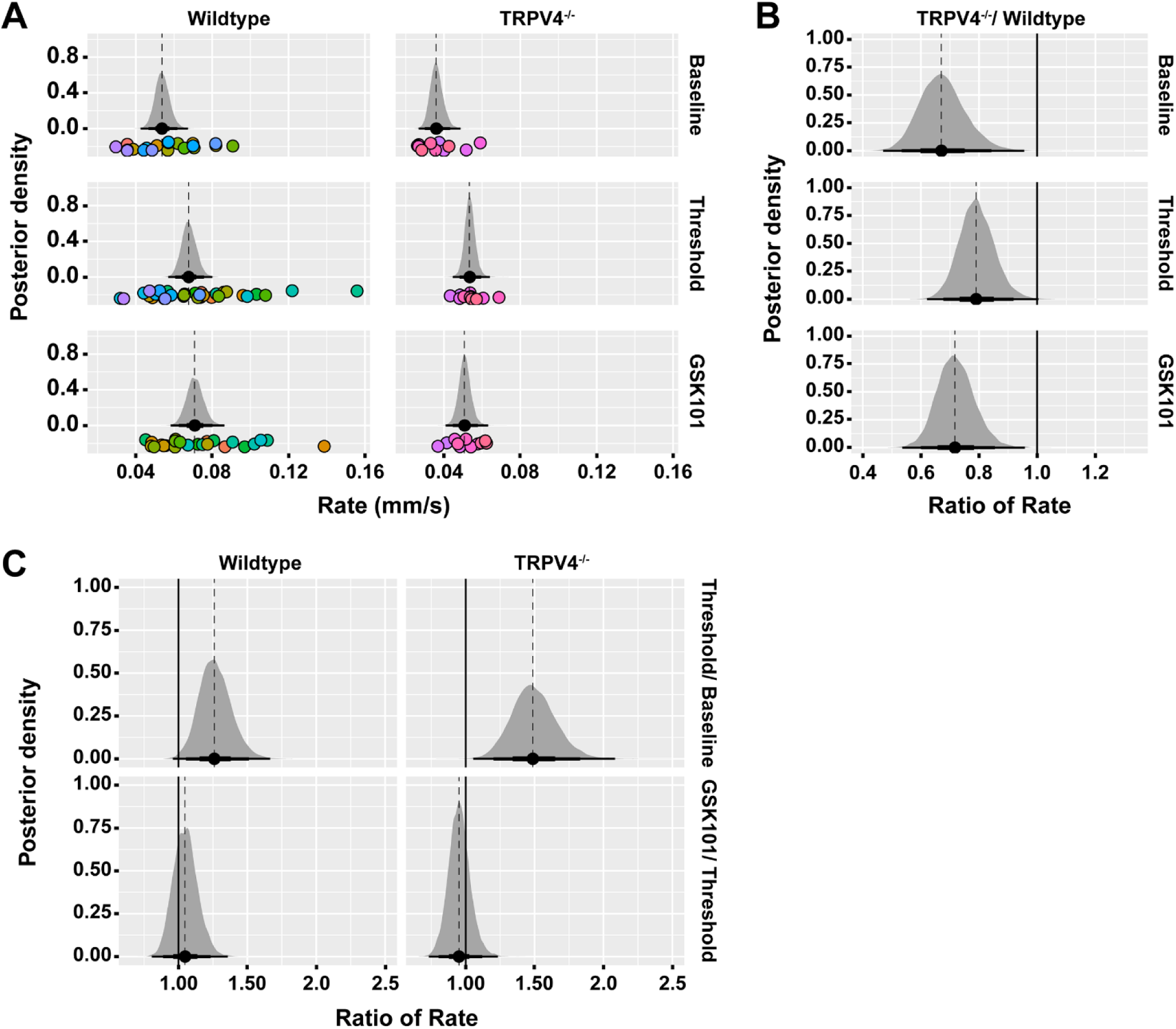
Deletion of TRPV4 reduces the mean rate of colonic contractions. **A.** Probability distribution graphs summarizing the MRC for wildtype (*left*) and TRPV4^-/-^ (*right*) colons under baseline, threshold, and GSK101-treated conditions (*top-bottom*). **B.** The ratio of MRCs was compared between wildtype and TRPV4^-/-^ colons at baseline (n=20 and n=11, *top*), threshold (n=34, n=11, *middle*) and GSK101-treated conditions (n=24 and n=11, *bottom*). The MRC was lower in TRPV4^-/-^ tissue compared to wildtype under all conditions tested (ratio of mean-baseline= 0.67, threshold= 0.90, GSK101= 0.89). **C.** Separate comparisons were made for the ratio of MRC at threshold/baseline (*top*) and threshold/GSK101 (*bottom*) for wildtype (*left*) and TRPV4^-/-^ colons (*right*), respectively. The MRC was greater at threshold compared to baseline for wildtype and TRPV4^-/-^ tissues (ratio of mean: wildtype= 1.11, TRPV4^-/-^= 1.48). There was no difference in the MRC at threshold when compared to GSK101-treated (ratio of mean: WT= 0.96, TRPV4^-/-^= 0.95). Each colored dot represents an individual experiment.

### Supplementary Videos

https://figshare.com/articles/media/Rajasekhar_et_al_2024_Supplemental_Videos/24956184

**Video S1. TRPV4 is functionally expressed by enteric glia and macrophages of the mouse colon.** Video shows Ca^2+^ imaging of a myenteric wholemount preparation of the mouse distal colon loaded with Fluo8-AM. TRPV4 activation (GSK101, 100nM, t=60s) evoked a rapid elevation in intracellular Ca^2+^ in mMac (CD68+, arrows) and enteric glia (Sox10+/GFAP+, circles). Responses by mMac were sustained, whereas Ca^2+^ oscillations were detected in glial cells. No responses were observed in neurons (Hu). Immunolabeling of the same ganglion is shown on the right.

**Video S2. Functional expression of TRPV4 in the human colon.** Video shows Ca^2+^ imaging of a submucosal wholemount preparation of the human colon. TRPV1 activation (capsaicin, 100nM) evoked a rapid elevation in Ca^2+^ in nerve fibers and varicosities (white arrows), which was followed by subsequent signaling in enteric neurons (magenta). Treatment with GSK101 (100nM) was associated with activation of enteric glial cells and vasculature (asterisks), but not nerve fibers (PGP9.5+) or neurons (HuC/D+).

**Video S3. TRPV4-mediated signaling in the glial network is retained with purinergic block.** Video shows Ca^2+^ imaging of a myenteric wholemount preparation of the mouse distal colon loaded with Fluo8-AM. GSK101 (100nM, 98s) evoked Ca^2+^ oscillations were retained in the presence of the purinoceptor antagonists suramin (50 µM) and PPADS (50 µM). Responses by glia to NKA (1µM, 592s) were retained under these conditions, but those to ATP (100µM, 660s) were abolished, confirming effective purinoceptor block.

**Video S4. The SERCA inhibitor cyclopiazonic acid (CPA; 30 µM) significantly enhanced responses by glial cells to GSK101.** Video shows Ca^2+^ imaging of a myenteric wholemount preparation of the mouse distal colon loaded with Fluo8-AM. Pretreatment with CPA increased the number of glial cells that were responsive to GSK101 (100nM, 69s). Effective block of SERCA was confirmed by the absence of Ca^2+^ responses to NKA (1µM, 429s). In contrast, there was no effect on responses to ATP (100µM, 488s), demonstrating that this was not due to a generalized inhibitory effect of CPA.

**Video S5. Oscillations in TRPV4-dependent Ca^2+^ signaling are via gap junctions.** Video shows Ca^2+^ imaging of a myenteric wholemount preparation of the mouse distal colon loaded with Fluo8-AM. Pretreatment with carbenoxolone (CBX, 50µM) suppressed Ca^2+^ oscillations and revealed a glial subset that exhibited rapid and sustained responses to GSK101 (100nM, 84s). Examples of sustained glial responses are indicated by arrows.

**Video S6. Acute inflammation is associated with a significant enhancement of glial responses to GSK101.**

Video shows imaging of a myenteric wholemount preparation of the acutely inflamed colon of a TNBS-treated mouse. TRPV4 activation (GSK101, 100nM, 62s) was associated with rapid elevation in Ca^2+^ in enteric glia, in addition to prominent activation of infiltrating immune cells.

**Video S7. Extrinsic afferent nerves respond to TRPV4 and TRPV1 activation.** Video shows a serosal afferent nerve in a wholemount preparation of the Wnt1-GCaMP3 mouse colon. Weak responses by this nerve to GSK101 (100nM, 37s) were followed by robust elevations in Ca^2+^ to capsaicin (500nM, 350s).

## Notes

https://figshare.com/articles/media/Rajasekhar_et_al_2024_Supplemental_Videos/24956184

### Competing Interest Statement

The authors have declared no competing interest.

## REFERENCES

1 Spencer NJ, Zagorodnyuk V, Brookes SJ, Hibberd T. Spinal afferent nerve endings in visceral organs: recent advances. Am J Physiol Gastrointest Liver Physiol 2016;311:G1056–G63.

2 Treichel AJ, Finholm I, Knutson KR, Alcaino C, Whiteman ST, Brown MR, et al. Specialized Mechanosensory Epithelial Cells in Mouse Gut Intrinsic Tactile Sensitivity. Gastroenterology 2022;162:535–47 e13.

3 Servin-Vences MR, Lam RM, Koolen A, Wang Y, Saade DN, Loud M, et al. PIEZO2 in somatosensory neurons controls gastrointestinal transit. Cell 2023;186:3386–99 e15.

4 Brierley SM, Page AJ, Hughes PA, Adam B, Liebregts T, Cooper NJ, et al. Selective role for TRPV4 ion channels in visceral sensory pathways. Gastroenterology 2008;134:2059–69.

5 Cenac N, Altier C, Chapman K, Liedtke W, Zamponi G, Vergnolle N. Transient receptor potential vanilloid-4 has a major role in visceral hypersensitivity symptoms. Gastroenterology 2008;135:937–46, 46 e1-2.

6 Peng S, Grace MS, Gondin AB, Retamal JS, Dill L, Darby W, et al. The transient receptor potential vanilloid 4 (TRPV4) ion channel mediates protease activated receptor 1 (PAR1)-induced vascular hyperpermeability. Lab Invest 2020;100:1057–67.

7 Swain SM, Romac JM, Shahid RA, Pandol SJ, Liedtke W, Vigna SR, et al. TRPV4 channel opening mediates pressure-induced pancreatitis initiated by Piezo1 activation. J Clin Invest 2020;130:2527–41.

8 Veldhuis NA, Poole DP, Grace M, McIntyre P, Bunnett NW. The G protein-coupled receptor-transient receptor potential channel axis: molecular insights for targeting disorders of sensation and inflammation. Pharmacol Rev 2015;67:36–73.

9 White JP, Cibelli M, Urban L, Nilius B, McGeown JG, Nagy I. TRPV4: Molecular Conductor of a Diverse Orchestra. Physiol Rev 2016;96:911–73.

10 Poole DP, Lieu T, Veldhuis N, Rajasekhar P, Bunnett NW. Targeting of Transient Receptor Potential Channels in Digestive Disease. In: Szallasi A, ed. TRP Channels as Therapeutic Targets From Basic Science to Clinical Use: Academic Press, 2015:385-403.

11 D’Aldebert E, Cenac N, Rousset P, Martin L, Rolland C, Chapman K, et al. Transient receptor potential vanilloid 4 activated inflammatory signals by intestinal epithelial cells and colitis in mice. Gastroenterology 2011;140:275–85.

12 Fichna J, Mokrowiecka A, Cygankiewicz AI, Zakrzewski PK, Malecka-Panas E, Janecka A, et al. Transient receptor potential vanilloid 4 blockade protects against experimental colitis in mice: a new strategy for inflammatory bowel diseases treatment? Neurogastroenterol Motil 2012;24:e557–60.

13 Matsumoto K, Yamaba R, Inoue K, Utsumi D, Tsukahara T, Amagase K, et al. Transient receptor potential vanilloid 4 channel regulates vascular endothelial permeability during colonic inflammation in dextran sulphate sodium-induced murine colitis. Br J Pharmacol 2018;175:84–99.

14 Cenac N, Bautzova T, Le Faouder P, Veldhuis NA, Poole DP, Rolland C, et al. Quantification and Potential Functions of Endogenous Agonists of Transient Receptor Potential Channels in Patients With Irritable Bowel Syndrome. Gastroenterology 2015;149:433–44 e7.

15 McGuire C, Boundouki G, Hockley JRF, Reed D, Cibert-Goton V, Peiris M, et al. Ex vivo study of human visceral nociceptors. Gut 2018;67:86–96.

16 Sipe WE, Brierley SM, Martin CM, Phillis BD, Cruz FB, Grady EF, et al. Transient receptor potential vanilloid 4 mediates protease activated receptor 2-induced sensitization of colonic afferent nerves and visceral hyperalgesia. Am J Physiol Gastrointest Liver Physiol 2008;294:G1288–98.

17 Fichna J, Poole DP, Veldhuis N, MacEachern SJ, Saur D, Zakrzewski PK, et al. Transient receptor potential vanilloid 4 inhibits mouse colonic motility by activating NO-dependent enteric neurotransmission. J Mol Med (Berl) 2015;93:1297–309.

18 Balemans D, Aguilera-Lizarraga J, Florens MV, Jain P, Denadai-Souza A, Viola MF, et al. Histamine-mediated potentiation of transient receptor potential (TRP) ankyrin 1 and TRP vanilloid 4 signaling in submucosal neurons in patients with irritable bowel syndrome. Am J Physiol Gastrointest Liver Physiol 2019;316:G338–G49.

19 Luo J, Qian A, Oetjen LK, Yu W, Yang P, Feng J, et al. TRPV4 Channel Signaling in Macrophages Promotes Gastrointestinal Motility via Direct Effects on Smooth Muscle Cells. Immunity 2018;49:107–19 e4.

20 Kollmann P, Elfers K, Maurer S, Klingenspor M, Schemann M, Mazzuoli-Weber G. Submucosal enteric neurons of the cavine distal colon are sensitive to hypoosmolar stimuli. J Physiol 2020;598:5317–32.

21 Sonkusare SK, Bonev AD, Ledoux J, Liedtke W, Kotlikoff MI, Heppner TJ, et al. Elementary Ca2+ signals through endothelial TRPV4 channels regulate vascular function. Science 2012;336:597–601.

22 Liedtke W, Friedman JM. Abnormal osmotic regulation in trpv4-/- mice. Proc Natl Acad Sci U S A 2003;100:13698–703.

23 Boesmans W, Martens MA, Weltens N, Hao MM, Tack J, Cirillo C, et al. Imaging neuron-glia interactions in the enteric nervous system. Front Cell Neurosci 2013;7:183.

24 Sasmono RT, Oceandy D, Pollard JW, Tong W, Pavli P, Wainwright BJ, et al. A macrophage colony-stimulating factor receptor-green fluorescent protein transgene is expressed throughout the mononuclear phagocyte system of the mouse. Blood 2003;101:1155–63.

25 Spencer NJ, Sorensen J, Travis L, Wiklendt L, Costa M, Hibberd T. Imaging activation of peptidergic spinal afferent varicosities within visceral organs using novel CGRPalpha-mCherry reporter mice. Am J Physiol Gastrointest Liver Physiol 2016;311:G880–G94.

26 Li Z, Khanna M, Grimley SL, Ellenberg P, Gonelli CA, Lee WS, et al. Mucosal IL-4R antagonist HIV vaccination with SOSIP-gp140 booster can induce high-quality cytotoxic CD4(+)/CD8(+) T cells and humoral responses in macaques. Sci Rep 2020;10:22077.

27 DiCello JJ, Rajasekhar P, Eriksson EM, Saito A, Gondin AB, Veldhuis NA, et al. Clathrin and GRK2/3 inhibitors block delta-opioid receptor internalization in myenteric neurons and inhibit neuromuscular transmission in the mouse colon. Am J Physiol Gastrointest Liver Physiol 2019;317:G79–G89.

28 Gavagnin E, Owen JP, Yates CA. Pair correlation functions for identifying spatial correlation in discrete domains. Phys Rev E 2018;97:062104.

29 Johnston ST, Crampin EJ. Corrected pair correlation functions for environments with obstacles. Phys Rev E 2019;99:032124.

30 Louis C, Cook AD, Lacey D, Fleetwood AJ, Vlahos R, Anderson GP, et al. Specific Contributions of CSF-1 and GM-CSF to the Dynamics of the Mononuclear Phagocyte System. J Immunol 2015;195:134–44.

31 Wirtz S, Popp V, Kindermann M, Gerlach K, Weigmann B, Fichtner-Feigl S, et al. Chemically induced mouse models of acute and chronic intestinal inflammation. Nat Protoc 2017;12:1295–309.

32 DiCello JJ, Carbone SE, Saito A, Pham V, Szymaszkiewicz A, Gondin AB, et al. Positive allosteric modulation of endogenous delta opioid receptor signaling in the enteric nervous system is a potential treatment for gastrointestinal motility disorders. Am J Physiol Gastrointest Liver Physiol 2022;322:G66–G78.

33 DiCello JJ, Saito A, Rajasekhar P, Sebastian BW, McQuade RM, Gondin AB, et al. Agonist-dependent development of delta opioid receptor tolerance in the colon. Cell Mol Life Sci 2019;76:3033–50.

34 Seguella L, Gulbransen BD. Enteric glial biology, intercellular signalling and roles in gastrointestinal disease. Nat Rev Gastroenterol Hepatol 2021;18:571–87.

35 Sharkey KA, Mawe GM. The enteric nervous system. Physiol Rev 2023;103:1487–564.

36 Retamal JS, Grace MS, Dill LK, Ramirez-Garcia P, Peng S, Gondin AB, et al. Serotonin-induced vascular permeability is mediated by transient receptor potential vanilloid 4 in the airways and upper gastrointestinal tract of mice. Lab Invest 2021;101:851–64.

37 Vergnolle N, Cenac N, Altier C, Cellars L, Chapman K, Zamponi GW, et al. A role for transient receptor potential vanilloid 4 in tonicity-induced neurogenic inflammation. Br J Pharmacol 2010;159:1161–73.

38 Shibasaki K, Ikenaka K, Tamalu F, Tominaga M, Ishizaki Y. A novel subtype of astrocytes expressing TRPV4 (transient receptor potential vanilloid 4) regulates neuronal excitability via release of gliotransmitters. J Biol Chem 2014;289:14470–80.

39 Dunn KM, Hill-Eubanks DC, Liedtke WB, Nelson MT. TRPV4 channels stimulate Ca2+-induced Ca2+ release in astrocytic endfeet and amplify neurovascular coupling responses. Proc Natl Acad Sci U S A 2013;110:6157–62.

40 Delvalle NM, Dharshika C, Morales-Soto W, Fried DE, Gaudette L, Gulbransen BD. Communication Between Enteric Neurons, Glia, and Nociceptors Underlies the Effects of Tachykinins on Neuroinflammation. Cell Mol Gastroenterol Hepatol 2018;6:321–44.

41 McClain JL, Fried DE, Gulbransen BD. Agonist-evoked Ca(2+) signaling in enteric glia drives neural programs that regulate intestinal motility in mice. Cell Mol Gastroenterol Hepatol 2015;1:631–45.

42 Zhang W, Segura BJ, Lin TR, Hu Y, Mulholland MW. Intercellular calcium waves in cultured enteric glia from neonatal guinea pig. Glia 2003;42:252–62.

43 Boesmans W, Hao MM, Fung C, Li Z, Van den Haute C, Tack J, et al. Structurally defined signaling in neuro-glia units in the enteric nervous system. Glia 2019;67:1167–78.

44 McClain J, Grubisic V, Fried D, Gomez-Suarez RA, Leinninger GM, Sevigny J, et al. Ca2+ responses in enteric glia are mediated by connexin-43 hemichannels and modulate colonic transit in mice. Gastroenterology 2014;146:497–507 e1.

45 Muller PA, Koscso B, Rajani GM, Stevanovic K, Berres ML, Hashimoto D, et al. Crosstalk between Muscularis Macrophages and Enteric Neurons Regulates Gastrointestinal Motility. Cell 2014;158:1210.

46 Grubisic V, McClain JL, Fried DE, Grants I, Rajasekhar P, Csizmadia E, et al. Enteric Glia Modulate Macrophage Phenotype and Visceral Sensitivity following Inflammation. Cell Rep 2020;32:108100.

47 Vergnolle N. TRPV4: new therapeutic target for inflammatory bowel diseases. Biochem Pharmacol 2014;89:157–61.

48 Drokhlyansky E, Smillie CS, Van Wittenberghe N, Ericsson M, Griffin GK, Eraslan G, et al. The Human and Mouse Enteric Nervous System at Single-Cell Resolution. Cell 2020;182:1606–22 e23.

49 Matheis F, Muller PA, Graves CL, Gabanyi I, Kerner ZJ, Costa-Borges D, et al. Adrenergic Signaling in Muscularis Macrophages Limits Infection-Induced Neuronal Loss. Cell 2020;180:64–78 e16.

50 Boesmans W, Lasrado R, Vanden Berghe P, Pachnis V. Heterogeneity and phenotypic plasticity of glial cells in the mammalian enteric nervous system. Glia 2015;63:229–41.

51 Rao M, Nelms BD, Dong L, Salinas-Rios V, Rutlin M, Gershon MD, et al. Enteric glia express proteolipid protein 1 and are a transcriptionally unique population of glia in the mammalian nervous system. Glia 2015;63:2040–57.

52 Seguella L, McClain JL, Esposito G, Gulbransen BD. Functional Intraregional and Interregional Heterogeneity between Myenteric Glial Cells of the Colon and Duodenum in Mice. J Neurosci 2022;42:8694–708.

53 Mapps AA, Thomsen MB, Boehm E, Zhao H, Hattar S, Kuruvilla R. Diversity of satellite glia in sympathetic and sensory ganglia. Cell Rep 2022;38:110328.

54 Turovsky EA, Braga A, Yu Y, Esteras N, Korsak A, Theparambil SM, et al. Mechanosensory Signaling in Astrocytes. J Neurosci 2020;40:9364–71.

55 Rao M, Rastelli D, Dong L, Chiu S, Setlik W, Gershon MD, et al. Enteric Glia Regulate Gastrointestinal Motility but Are Not Required for Maintenance of the Epithelium in Mice. Gastroenterology 2017;153:1068–81 e7.

56 Poole DP, Amadesi S, Veldhuis NA, Abogadie FC, Lieu T, Darby W, et al. Protease-activated receptor 2 (PAR2) protein and transient receptor potential vanilloid 4 (TRPV4) protein coupling is required for sustained inflammatory signaling. J Biol Chem 2013;288:5790–802.

57 Lucchinetti CF, Kimmel DW, Lennon VA. Paraneoplastic and oncologic profiles of patients seropositive for type 1 antineuronal nuclear autoantibodies. Neurology 1998;50:652–7.

## References

1 Rueden CT, Schindelin J, Hiner MC, DeZonia BE, Walter AE, Arena ET, et al. ImageJ2: ImageJ for the next generation of scientific image data. BMC Bioinformatics 2017;18:529.

2 Schneider CA, Rasband WS, Eliceiri KW. NIH Image to ImageJ: 25 years of image analysis. Nat Methods 2012;9:671–5.

3 Lowe DG. Distinctive Image Features from Scale-Invariant Keypoints. International Journal of Computer Vision 2004;60:91–110.

4 Arganda-Carreras I, Sorzano COS, Marabini R, Carazo JM, Ortiz-de-Solorzano C, Kybic J. Consistent and Elastic Registration of Histological Sections using Vector-Spline Regularization. Lecture Notes in Computer Science 2006;4241:85–95.

5 Gavagnin E, Owen JP, Yates CA. Pair correlation functions for identifying spatial correlation in discrete domains. Phys Rev E 2018;97:062104.

6 Johnston ST, Crampin EJ. Corrected pair correlation functions for environments with obstacles. Phys Rev E 2019;99:032124.

7 Ruhl A, Trotter J, Stremmel W. Isolation of enteric glia and establishment of transformed enteroglial cell lines from the myenteric plexus of adult rat. Neurogastroenterol Motil 2001;13:95–106.

8 Sampaio NG, Emery SJ, Garnham AL, Tan QY, Sisquella X, Pimentel MA, et al. Extracellular vesicles from early stage Plasmodium falciparum-infected red blood cells contain PfEMP1 and induce transcriptional changes in human monocytes. Cell Microbiol 2018;20:e12822.

9 de Almeida MC, Silva AC, Barral A, Barral Netto M. A simple method for human peripheral blood monocyte isolation. Mem Inst Oswaldo Cruz 2000;95:221–3.

10 Achuthan A, Cook AD, Lee MC, Saleh R, Khiew HW, Chang MW, et al. Granulocyte macrophage colony-stimulating factor induces CCL17 production via IRF4 to mediate inflammation. J Clin Invest 2016;126:3453–66.

11 Fichna J, Poole DP, Veldhuis N, MacEachern SJ, Saur D, Zakrzewski PK, et al. Transient receptor potential vanilloid 4 inhibits mouse colonic motility by activating NO-dependent enteric neurotransmission. J Mol Med (Berl) 2015;93:1297–309.

12 Rajasekhar P, Poole DP, Liedtke W, Bunnett NW, Veldhuis NA. P2Y1 Receptor Activation of the TRPV4 Ion Channel Enhances Purinergic Signaling in Satellite Glial Cells. J Biol Chem 2015;290:29051–62.

13 Louis C, Cook AD, Lacey D, Fleetwood AJ, Vlahos R, Anderson GP, et al. Specific Contributions of CSF-1 and GM-CSF to the Dynamics of the Mononuclear Phagocyte System. J Immunol 2015;195:134–44.

14 Wirtz S, Popp V, Kindermann M, Gerlach K, Weigmann B, Fichtner-Feigl S, et al. Chemically induced mouse models of acute and chronic intestinal inflammation. Nat Protoc 2017;12:1295–309.

15 DiCello JJ, Rajasekhar P, Eriksson EM, Saito A, Gondin AB, Veldhuis NA, et al. Clathrin and GRK2/3 inhibitors block delta-opioid receptor internalization in myenteric neurons and inhibit neuromuscular transmission in the mouse colon. Am J Physiol Gastrointest Liver Physiol 2019;317:G79–G89.

16 DiCello JJ, Saito A, Rajasekhar P, Sebastian BW, McQuade RM, Gondin AB, et al. Agonist-dependent development of delta opioid receptor tolerance in the colon. Cell Mol Life Sci 2019;76:3033–50.

17 DiCello JJ, Carbone SE, Saito A, Pham V, Szymaszkiewicz A, Gondin AB, et al. Positive allosteric modulation of endogenous delta opioid receptor signaling in the enteric nervous system is a potential treatment for gastrointestinal motility disorders. Am J Physiol Gastrointest Liver Physiol 2022;322:G66–G78.

18 Costa M, Wiklendt L, Arkwright JW, Spencer NJ, Omari T, Brookes SJ, et al. An experimental method to identify neurogenic and myogenic active mechanical states of intestinal motility. Front Syst Neurosci 2013;7:7.

19 Burkner P-C. brms: An R Package for Bayesian Multilevel Models Using Stan. Journal of Statistical Software 2017;80:1–28.

